# Collagen type I promotes pancreatic tumor growth and limits immune cell infiltration

**DOI:** 10.64898/2026.01.09.698616

**Authors:** Marie-Louise Thorseth, Astrid Z. Johansen, Kevin J. Baker, Marco Carretta, Anne Mette Askehøj Rømer, Christina Jensen, Henrik J. Jürgensen, Hannes Linder, Nadia K. Czajkowski, Dorota E. Kuczek, Klaire Y. Fjæstad, Lars H. Engelholm, Nicholas Willumsen, Kevin Kim, Mads H. Andersen, Lars Grøntved, Daniel H. Madsen

**Affiliations:** National Center for Cancer Immune Therapy (CCIT-DK), Department of Oncology, Copenhagen University Hospital – Herlev and Gentofte, Herlev, Denmark; Biomarkers and Research, Nordic Bioscience, Herlev, Denmark; Finsen Laboratory, Rigshospitalet/Biotech Research and Innovation Centre, University of Copenhagen, Copenhagen, Denmark; Department of Internal Medicine, University of Michigan, Ann Arbor, USA; Department of Immunology and Microbiology, University of Copenhagen, Copenhagen, Denmark; Department of Biochemistry and Molecular Biology, University of Southern Denmark, Odense, Denmark

**Keywords:** Pancreatic cancer, Extracellular matrix remodeling, Tumor immune microenvironment, Cancer-associated fibrosis, Natural killer cell infiltration, Myeloid-derived suppressor cells, Tumor–stroma interactions

## Abstract

Solid tumors are often characterized by a dense extracellular matrix (ECM) that contributes to increased tissue stiffness. Collagen type I is the main component of the ECM and its abundance in tumors is frequently associated with poor prognosis. In vitro studies suggest that a high collagen density promotes tumor invasion and modulates immune responses. However, recent in vivo findings have questioned the pro-tumorigenic role of collagen type I.

In this study, we investigate the role of collagen for pancreatic tumor growth and immune cell infiltration using conditional collagen type I knockout mice and transgenic collagenase-resistant mice. Preventing collagen type I significantly reduces intratumoral collagen content and tumor growth. This reduction is accompanied increased infiltration of natural killer (NK) cells, a higher CD8/CD4 T cell ratio, and decreased numbers of monocytic myeloid-derived suppressor cells (MDSCs).

Conversely, collagenase-resistant mice develop collagen-dense tumors and display enhanced tumor growth. These mice also generally exhibit opposing effects on cell infiltration, including a lower CD8/CD4 ratio and increased MDSC abundance.

These findings are further supported by analyses of publicly available human cancer datasets, which confirm an association between collagen type I levels and immune cell infiltration.

Overall, our results demonstrate a pronounced pro-tumorigenic role of collagen type I in pancreatic cancer, which is associated with modulation of the tumor immune microenvironment. This study highlights the importance of extracellular matrix components as key regulators of tumor progression and anti-tumor immunity.

## Introduction

Solid tumors are complex tissues composed not only of malignant cells but also of stromal and immune cells embedded within a dynamic extracellular matrix (ECM) ^1^. A hallmark of many solid tumors is the extensive remodeling of the ECM, including increased deposition and reorganization of collagen type I (Col1) ^2–4^. During tumor progression, Col1 accumulation results in a dense, fibrotic ECM that is markedly different from the ECM of normal tissues. The tumor-specific ECM is typically stiffer and organized in aligned fibers ^2,3,5^, and these changes have been associated with enhanced tumor aggressiveness and invasiveness ^1,6,7^.

Col1 is the main component of the ECM and the most abundant protein in the mammalian body. Col1 is a heterotrimer composed of two α1(I) chains and one α2(I) chain that form a tight triple-helical structure ^8,9^. Upon secretion, Col1 molecules self-assemble into insoluble fibrils that are further stabilized through enzymatic crosslinking, predominantly mediated by lysyl oxidase (LOX) ^10^. Due to its unique structure, Col1 is highly resistant to proteolytic degradation ^11^. However, a limited number of matrix metalloproteinases (MMPs), such as MMP-1, MMP-13, and MMP-14 (MT1-MMP), can cleave Col1 at specific sites, yielding well-defined ¾ and ¼ fragments ^12–14^. These fragments can be further processed by gelatinases or internalized by stromal cells such as cancer-associated fibroblasts (CAFs) and tumor-associated macrophages (TAMs) for lysosomal degradation ^15–20^. These turnover dynamics contribute to the ECM remodeling observed during cancer progression.

Numerous clinical and preclinical studies have shown that increased collagen density within tumors correlates with poor prognosis in several cancer types, including breast, gastric, pancreatic, and oral squamous cell carcinomas ^21–25^. However, a direct causal link between collagen density and poor prognosis has not been clearly established. Studies in mice have indicated that the collagen density can directly affect breast cancer growth. Specifically, increased collagen density of tumors grown in transgenic Col1a1^tm1jae^ (Col^r^) mice has been associated with faster cancer growth and enhanced metastasis ^26–29^. Col^r^ mice harbor mutations in the collagenase cleavage site of the collagen I α1-chain, which results in strongly impaired collagen turnover and increased collagen accumulation ^26,28,30^. Furthermore, high-density Col1 matrices have been shown to promote malignant transformation, stimulate epithelial-to-mesenchymal transition (EMT), and enhance cancer cell invasion and migration *in vitro* ^26,31^. Overall, the ECM can influence several of the hallmarks of cancer, such as sustaining proliferative signaling, inducing angiogenesis, and activating invasion and metastasis ^1,32^.

The ECM can also affect immune cell activation and infiltration into tumors ^6,33,34^. Specifically, Col1 has been shown to dictate the migratory trajectory of T cells and limit their infiltration into the tumors ^35,36^. We have previously demonstrated that high-density Col1 matrices can also directly suppress T cell cytotoxicity and promote immunosuppressive polarization of macrophages in *ex vivo* 3D culture models ^37,38^. In pancreatic cancer, aligned collagen fibers have recently been shown to associate with an immunosuppressive macrophage phenotype, which in turn can limit T cell activity and migration ^39^.

However, the view of collagen as an exclusively pro-tumorigenic entity has recently been challenged. Two independent studies using cell type-specific deletion of Col1 in mouse models of pancreatic and metastatic liver cancer found that reduced collagen deposition was associated with increased tumor aggressiveness and reduced overall survival ^40,41^. These surprising findings suggest that the role of Col1 in cancer is complex and context-dependent and that it may involve tumor-suppressive functions in specific settings.

In the present study, we sought to clarify the role of collagen in tumor progression and immune regulation by employing both loss-of-function and gain-of-function genetic mouse models. Using an inducible systemic Col1a1 knockout (KO) model, we show that decreased intratumoral collagen content leads to reduced tumor growth of subcutaneous pancreatic Pan02 tumors, possibly mediated by an increase in tumor-infiltrating immune effector cells and a decrease in immunosuppressive cells. To support these findings, we used the collagen-accumulating transgenic Col^r^ mouse model, in which we saw increased tumor growth, reduced infiltration of immune effector cells and increased infiltration of immunosuppressive cells, further supporting the pro-tumorigenic effects of collagen. Our findings provide new insights into the dualistic role of collagen in cancer and highlight the importance of ECM remodeling in shaping the immune landscape of solid tumors.

## Results

### Generation of a mouse model with systemic deletion of Col1a1

To study the role of Col1 in pancreatic cancer, we generated inducible Col1a1 KO mice. The previously described Col1a1^flox/flox^ mice have loxP sites flanking exons two to five of the *Col1a1* gene ^42^, which upon Cre recombinase-mediated excision leads to a frameshift, early termination of translation, truncated mRNA and non-functional α1-chains. In the absence of the α1-chains, Col1 fibers cannot form. In this study, Col1a1^flox/flox^ mice were interbred with β-actin-cre/ER mice, which have ubiquitous expression of a Cre recombinase fusion protein that is activated by interaction with tamoxifen. This breeding program in turn generated either double-transgenic β-actin-cre/ER;Col1a1^flox/flox^ (termed Col1a1 KO) or single-transgenic Col1a1^flox/flox^ (control) littermate offspring (fig. 1A).

**Figure 1.**
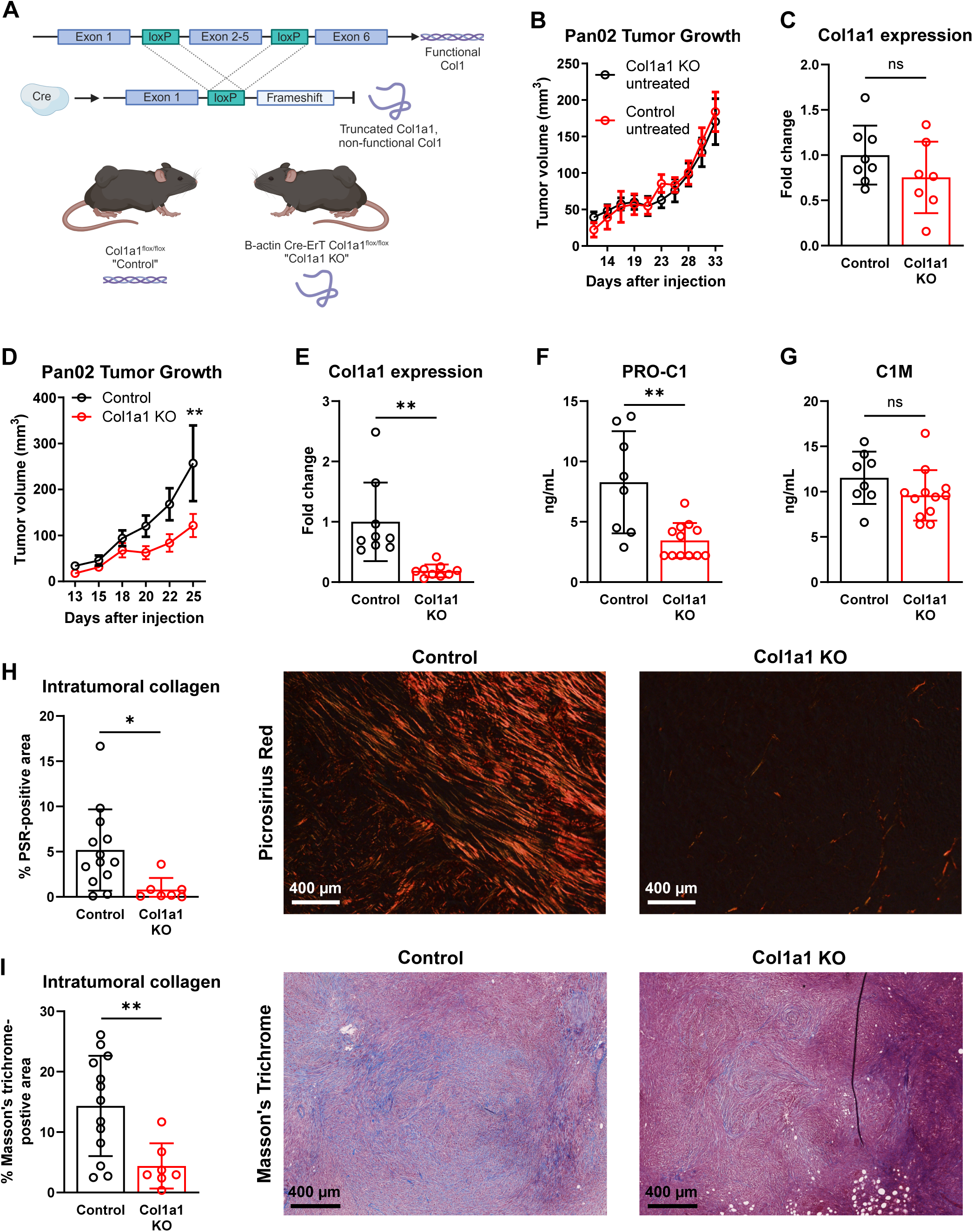
Systemic deletion of collagen type I decreases collagen deposition and tumor growth in a murine model of pancreatic cancer. **(A)** Schematic representation of the tamoxifen-inducible Col1a1 KO mouse model. Created with Biorender.com. **(B)** Mean tumor growth curves of wildtype littermate control and non-tamoxifen-treated Col1a1 KO mice. **(C)** *Col1a1* gene expression in tumors from control and non-tamoxifen-treated Col1a1 KO mice. **(B-C)** n = 7-8 mice per group. **(D)** Mean tumor growth curves of tamoxifen-treated control and Col1a1 KO mice. **(E)** *Col1a1* gene expression in tumors from tamoxifen-treated control and Col1a1 KO mice. **(D-E)** n = 8-9 mice per group. **(F-G)** Serum levels of the collagen formation product PRO-C1 **(F)** and the collagen degradation product C1M **(G)** from tamoxifen-treated control and Col1a1 KO mice. Some samples were below the lower limit of detection (2.2ng/mL for PRO-C1 and 6.38 for C1M). **(H-I)** Quantification (left) and representative images (right) of picrosirius red (PSR) **(H)** and Masson’s Trichrome (I) staining of Pan02 tumors from tamoxifen-treated control and Col1a1 KO mice. Each data point represents the average of 10 random squares per image. Statistical significance was assessed using the two-tailed student’s t-test **(C, E-I)** and two-way ANOVA with Bonferroni correction **(D)**. Bars represent mean ± SEM **(B,D)** or SD **(C, E-I)**. \**p*<0.05, \*\**p*<0.01, \*\*\**p*<0.001.

To evaluate if the inducible Cre recombinase system was leaky, we first inoculated Col1a1 KO mice and control wildtype littermates subcutaneously with the murine pancreatic ductal adenocarcinoma (PDAC) cell line Pan02 without administration of tamoxifen. Pan02 cancer cells form highly desmoplastic tumors compared to many other murine cancer models ^40,43^. Tumor growth in the two groups of mice were indistinguishable from each other (fig. 1B) and qRT-PCR analysis of RNA isolated from whole tumors revealed similar levels of *Col1a1* expression, demonstrating that recombination and deletion of Col1 does not happen in the absence of tamoxifen (fig. 1C).

### Systemic knockout of collagen type 1 reduces tumor growth in a model of pancreatic cancer

Next, Col1a1 KO and control mice were inoculated with Pan02, and tamoxifen administration was started when tumors became palpable. Strikingly, Col1a1 KO mice displayed significantly reduced tumor growth at day 25 after inoculation (fig. 1D). qRT-PCR analysis of whole tumors revealed that tamoxifen-treatment led to an 81% reduction in *Col1a1* expression in Col1a1 KO mice compared to the tamoxifen-treated control mice (fig. 1E). Reduced production of Col1 was confirmed by measuring serum levels of the PRO-C1 epitope in a competitive ELISA (fig. 1F). PRO-C1 is generated from the pro-peptide of Col1 during synthesis and was significantly reduced in serum from Col1a1 KO mice. There was no indication of a change in the level of Col1 degradation as similar serum levels of the MMP-specific Col1 degradation product C1M was measured (fig. 1G). The decreased synthesis without a compensatory decrease in degradation suggests an overall decreased deposition of Col1. To confirm this, tumor tissue sections were stained with picrosirius red (PSR) (fig. 1H) or Masson’s trichrome stain (fig. 1I). Both types of staining revealed a significant reduction in deposited fibrillar collagen in tumors from Col1a1 KO mice compared to tumors from control mice. Phenotypically, Col1a1 KO mice could not be distinguished from wildtype littermates before initiation of tamoxifen administration. About 21 days after cancer cell inoculation and 11 days after tamoxifen-induced deletion of Col1, Col1a1 KO mice started to present with stiff movements and weight loss (fig. S1). In most cases, Col1a1 KO mice had to be euthanized within 4 weeks due to wasting. Wildtype littermates did not show any of the same symptoms.

### Col1a1 KO mice have more tumor-infiltrating lymphocytes

To investigate if the reduced collagen levels in the tumors impacted the abundance of tumor-infiltrating immune cells, tumors were excised, enzymatically digested to a single-cell suspension, and analyzed by multi-color flow cytometry using three different antibody panels (Table S1 and fig. S2). The overall cellular composition was similar between tumors from Col1a1 KO mice and wildtype littermates when comparing ratios of cancer cells (CD45^-^CD31^-^FAP^-^PDGFRα^-^), immune cells (CD45^+^), CAFs (CD45^-^CD31^-^FAP^+^PDGFRα^+^;CD45^-^CD31^-^FAP^+^PDGFRα^-^;CD45^-^CD31^-^FAP^-^PDGFRα+), and endothelial cells (CD45^-^CD31^+^) (fig. 2A). Expression of the immune inhibitory receptor PD-L1 ^44^ was slightly reduced on immune cells in Col1a1 KO tumors, although this difference was not statistically significant (fig. 2B). There was no difference in the expression level of PD-L1 on cancer cells but a trend towards reduced PD-L1 on CAFs (fig. 2C-D). The trends of decreased PD-L1 expression on the most abundant cell populations of the tumor (fig. 2B-D) was confirmed by qRT-PCR analysis of total tumor RNA showing significantly reduced gene expression levels of *Cd274* encoding PD-L1 in Col1a1 KO mice (fig. 2E).

**Figure 2.**
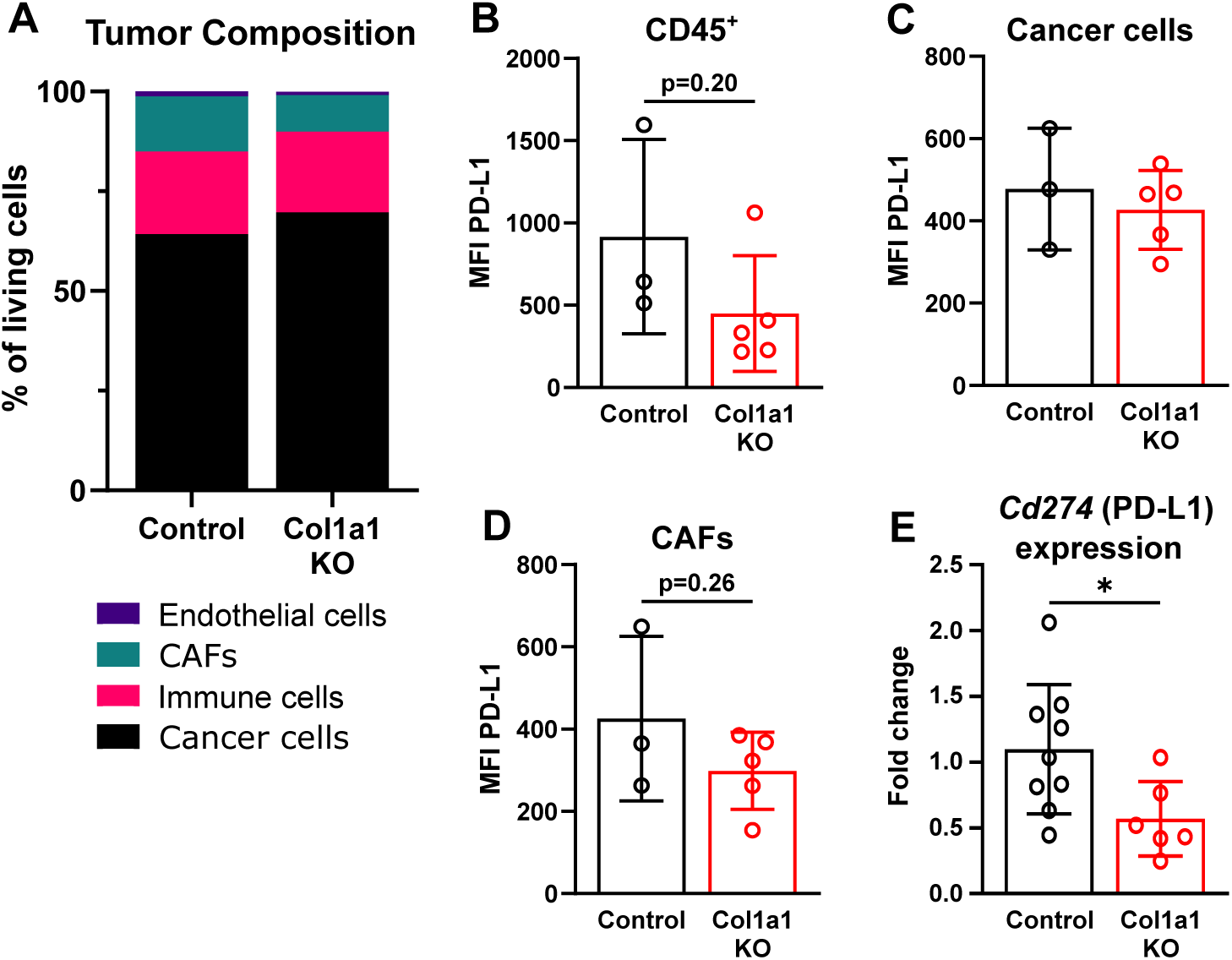
Collagen type I knockout alters PD-L1 expression in the tumor microenvironment. **(A)** Tumor composition quantified by flow cytometry: cancer cells (CD45^-^FAP^-^CD31^-^), immune cells (CD45^+^), CAFs (CD45^-^FAP^+^PDGFRα^+^; CD45^-^FAP^-^PDGFRα^+^; CD45^-^FAP^+^PDGFRα^-)^, and endothelial cells (CD45^-^CD31^+^) (n = 3-5 mice per group). **(B-D)** Mean fluorescent intensity (MFI) of PD-L1 on immune cells **(B)**, cancer cells **(C)**, and CAFs **(D)**. **(E)** *Cd274* gene expression in tumors from tamoxifen-treated control and Col1a1 KO mice. Statistical significance was assessed using the two-tailed student’s t-test. Bars represent mean ± SD. \**p*<0.05.

Collagen has previously been shown to affect lymphocyte infiltration into tumors ^35,36,45^ as well as their cytotoxic activity ^6,38^. Therefore, we investigated if reduced intratumoral collagen levels caused changes to the tumor-infiltrating lymphocyte population. Flow cytometry-based analysis of tumors from Col1a1 KO and control mice showed similar abundance of T cells (CD3^+^) (fig. 3A). However, trends toward fewer CD4^+^ T cells and more CD8^+^ T cells in the tumors of Col1a1 KO mice compared to control mice resulted in a significant reduction in the CD4/CD8T cell ratio (fig. 3B-D). B cells (CD19^+^) were not affected by KO of Col1 (fig. 3E) but, interestingly, the amount of NK cells was significantly increased in tumors from Cola1 KO mice (fig. 3F).

**Figure 3.**
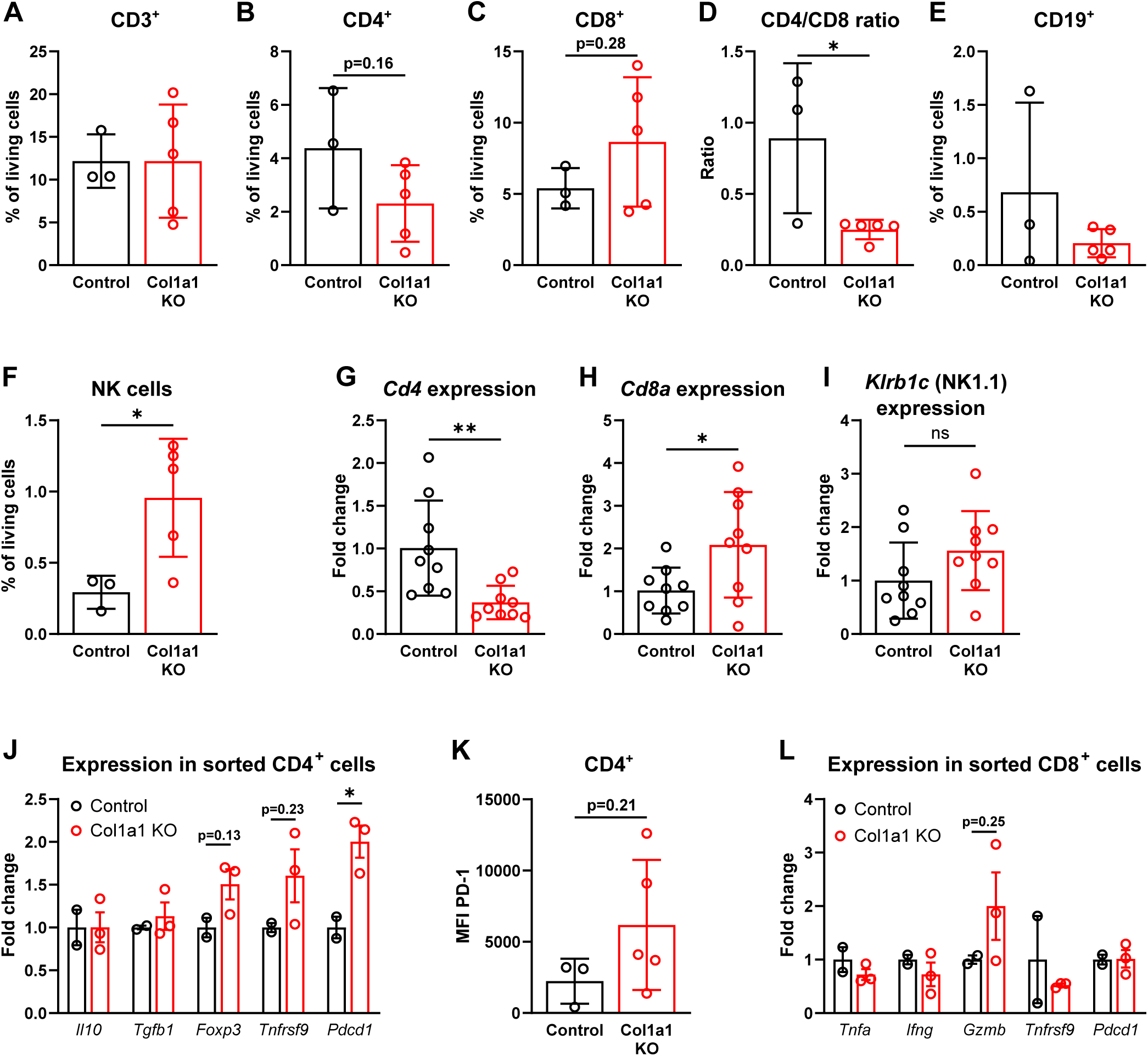
Collagen type I knockout affects lymphoid numbers and T cell phenotype. Lymphoid cell composition in Pan02 tumors quantified by flow cytometry: **(A)** T cells (CD3^+^), **(B)** CD4^+^ T cells, **(C)** CD8^+^ T cells, **(D)** CD4/CD8 ratio, **(E)** B cells (CD19^+^), and **(F)** NK cells (NK1.1^+^). **(G-I)** Gene expression of *Cd4* **(G)**, *Cd8a* **(H)** and *Klrb1c* **(I)** in tumors from tamoxifen-treated control and Col1a1 mice. **(J)** Gene expression of activation, exhaustion, and regulatory markers in FACS-sorted CD4^+^ T cell from tumors in tamoxifen-treated control and Col1a1 KO mice. **(K)** MFI of PD-1 in CD4^+^ T cells. **(L)** Gene expression of activation and exhaustion markers in FACS-sorted CD8^+^ T cells from tumors in tamoxifen-treated control and Col1a1 KO mice. Statistical significance was assessed using the two-tailed student’s t-test. Bars represent mean ± SD **(A-I, K)** or SEM **(J, L)**. \**p*<0.05, \*\**p*<0.01.

To validate these findings, qRT-PCR analyses of whole tumors were used to investigate the expression level of *Cd4*, *Cd8a,* and *Klrb1c* (encoding the NK cell marker NK1.1). The gene expression changes matched the flow cytometry data, since a 63% reduction in *Cd4* gene expression (fig. 3G) and a 105% increase in *Cd8a* gene expression (fig. 3H) was determined in tumors from Col1a1 KO mice compared to control. A 56% increase was recorded for *Klrb1c* gene expression although this difference did not achieve statistical significance (fig. 3I). To further explore if collagen affected the phenotype of tumor-infiltrating T cells, we used fluorescence-activated cell sorting (FACS) to isolate CD4^+^ and CD8^+^ T cells from tumors of Col1a1 KO mice and control wildtype littermates and analyzed gene expression levels of different markers of T cell activity (*Tnfrsf9, Tnfa, Ifng* and *Gzmb*), regulatory T cells (*Il10, Tgfb1* and *Foxp3*) and T cell exhaustion (*Pdcd1*) using qRT-PCR. In sorted CD4^+^ T cells from tumors from Col1a1 KO mice, there was a trend towards increased expression of *Foxp3* and *Tnfrsf9* and a significant increase in expression of *Pdcd1* encoding PD-1 (fig. 3J). A trend of increased PD1 was also apparent based on flow cytometry analysis (fig. 3K). In sorted CD8^+^ T cells from tumors from Col1a1 KO mice, there was a trend towards increased expression of *Gzmb* (fig. 3L).

Altogether, these results suggest that low collagen content in tumors favors the presence of cytotoxic lymphocytes, with a decreased CD4/CD8 ratio and higher abundance of NK cells, all of which can contribute to anti-tumor immunity.

### Col1a1 KO mice have fewer tumor-infiltrating myeloid-derived suppressor cells

Next, we evaluated the composition of myeloid cells in tumors from Col1a1 KO mice and control mice. The populations of analyzed myeloid cells included TAMs (CD11b^+^F4/80^+^), monocytic myeloid-derived suppressor cells (mMDSCs (CD11b^+^F4/80^-^CD11c^-^Ly-6C^+^Ly-6G^-^)), granulocytic myeloid-derived suppressor cells (gMDSCs (CD11b^+^F4/80^-^CD11c^-^Ly-6C^+^Ly-6G^+^)), and dendritic cells (DCs (CD11b^+^F4/80^-^CD11c^+^)). TAMs, mMDSCs and gMDSCs can promote an immunosuppressive tumor microenvironment (TME) and are associated with poor prognosis in many cancers ^46^. The abundance of TAMs was slightly, but not significantly, reduced in tumors from Col1a1 KO mice (fig. 4A), while the expression of the M2-like macrophage marker, MR, was not affected (fig. 4B). The number of mMDSCs was significantly reduced (Fig. 4C), whereas the number of gMDSCs was unaffected (Fig. 4D). These changes could indicate the formation of a less immunosuppressive TME in Col1a1 KO tumors.

**Figure 4.**
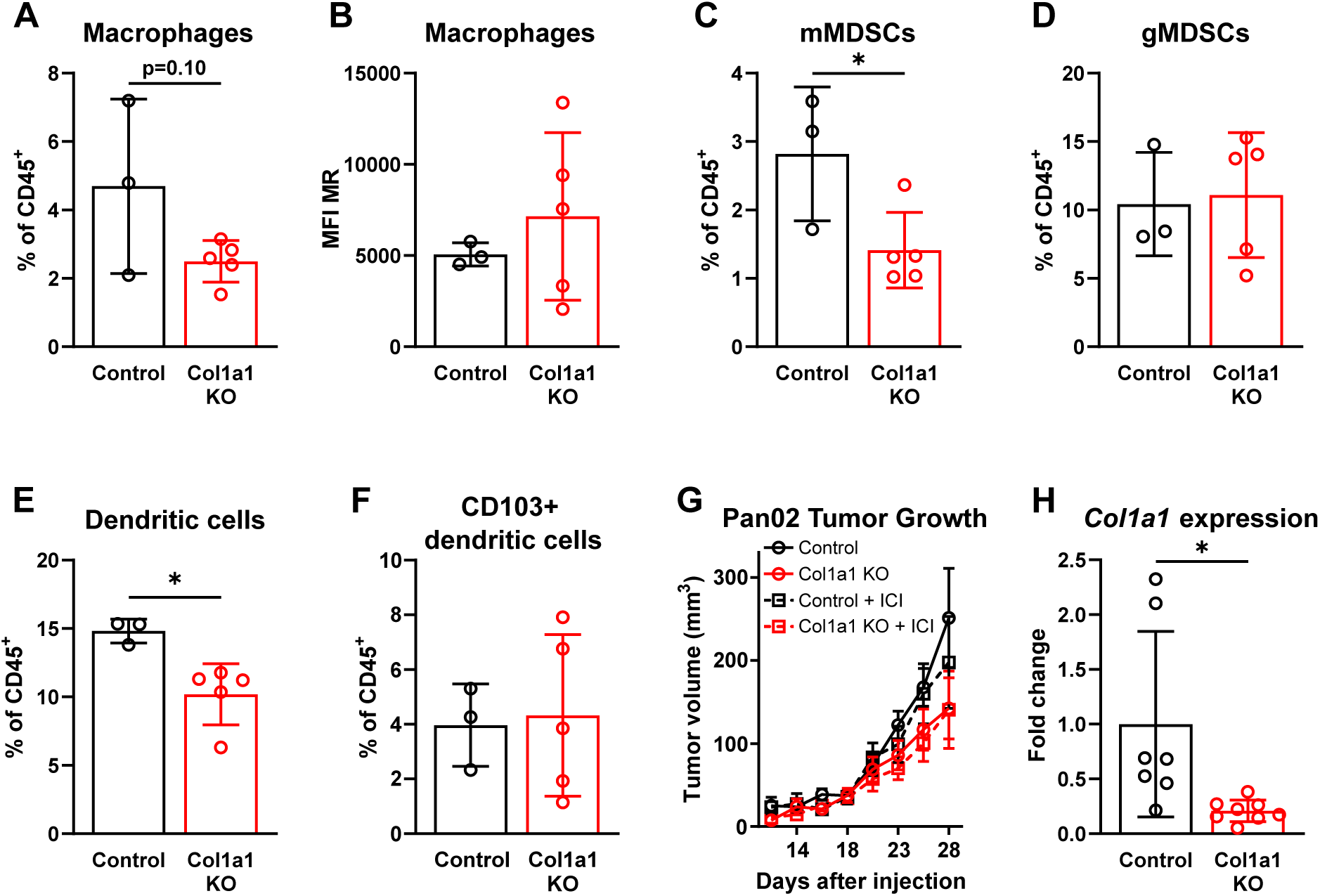
Collagen type I knockout reduces infiltration of immunosuppressive myeloid cells but does not enhance response to ICI therapy. Myeloid cell composition in Pan02 tumors from tamoxifen-treated control and Col1a1 KO mice quantified by flow cytometry: **(A)** macrophages (CD11b^+^F4/80^+^), **(B)** MFI of MR on macrophages, **(C)** mMDSCs (CD11b^+^F4/80^-^CD11c^-^Ly-6C^+^Ly-6G^-^), **(D)** gMDSCs (CD11b^+^F4/80^-^CD11c^-^Ly-6C^+^Ly-6G^+^), **(E)** dendritic cells (CD11b^+^F4/80^-^CD11c^+^), and **(F)** CD103^+^ dendritic cells. **(G)** *Col1a1* gene expression in tumors from tamoxifen-treated control and Col1a1 KO mice. **(H)** Mean Pan02 tumor growth of tamoxifen-treated control and Col1a1 KO mice, divided into control treated or ICI treatment with a combination of anti-CTLA-4 and anti-PD-L1 immune checkpoint inhibitors 10 days after tumor inoculation (n = 8-9 per group). Statistical significance was assessed using the two-tailed student’s t-test (A-G). Bars represent mean ± SD **(A-G)** or SEM **(H)**. \**p*<0.05.

The total amount of DCs was also reduced in tumors from Col1a1 KO mice (Fig. 4E) but the number of CD103^+^ DCs was unaltered (fig. 4F). The CD103^+^ DCs have been shown to represent a subset of DCs critical for eliciting an efficient anti-tumor immune response ^47,48^.

### Reduced intratumoral collagen type I does not improve sensitivity to immune checkpoint inhibitor therapy

Increased tumor-infiltration of immune effector cells and decreased infiltration of MDSCs are two factors that have been shown to correlate with increased response to immune checkpoint inhibitor (ICI) treatment ^46,49^. The Pan02 tumor model is non-immunogenic and unresponsive to ICI therapy ^50,51^, but the observed changes in the TME of tumors from Col1a1 KO mice led us to speculate if reduced collagen type I levels could sensitize the Pan02 tumors to ICI treatment. Col1a1 KO mice and control wildtype littermates were inoculated subcutaneously with Pan02 cells and administration of tamoxifen, anti-PD-L1 and anti-CTLA4 commenced when tumors became palpable. In this experiment we observed a similar reduction (79%) in *Col1a1* gene expression as previously observed (Fig. 4G, compare to Fig. 1E) and a similar effect of the Col1a1 KO on tumor growth (Fig. 4H). However, there was no additive effect of combining ICI therapy with Col1a1 gene inactivation (Fig. 4H), showing that a reduction in collagen type I levels was not sufficient for sensitizing the Pan02 tumors to ICI treatment.

### Increased level of collagen promotes the growth of Pan02 tumors

Using the Col1a1 KO mice, we demonstrated that a reduced level of collagen impedes pancreatic tumor growth and promotes the infiltration of cytotoxic immune cells. To evaluate if an enhanced collagen level would have opposite effects, we employed a different transgenic model, the Col^r^ mice ^30^. These mice harbor a mutation in the *Col1a1* gene rendering the collagen type I largely resistant to MMP-mediated degradation, which in turn leads to increased collagen accumulation. The mice were subcutaneously inoculated with Pan02 cells and tumor growth was measured. Collagen accumulation in the Col^r^ mice led to increased tumor growth compared to wildtype littermates (fig. 5A). Flow-cytometry based analyses showed that collagen accumulation caused a significant decrease in the abundance of tumor-infiltrating CD45^+^ immune cells and a corresponding increase in cancer cells (fig. 5B). In Col^r^ mice there was a trend of increased PD-L1 expression on immune cells (fig. 5C), whereas PD-L1 expression on CAFs and cancer cells was unaffected (fig. 5D-E). Analysis of total tumor RNA also showed increased *Cd274* gene expression levels in tumors from Col^r^ mice (fig. 5F).

**Figure 5.**
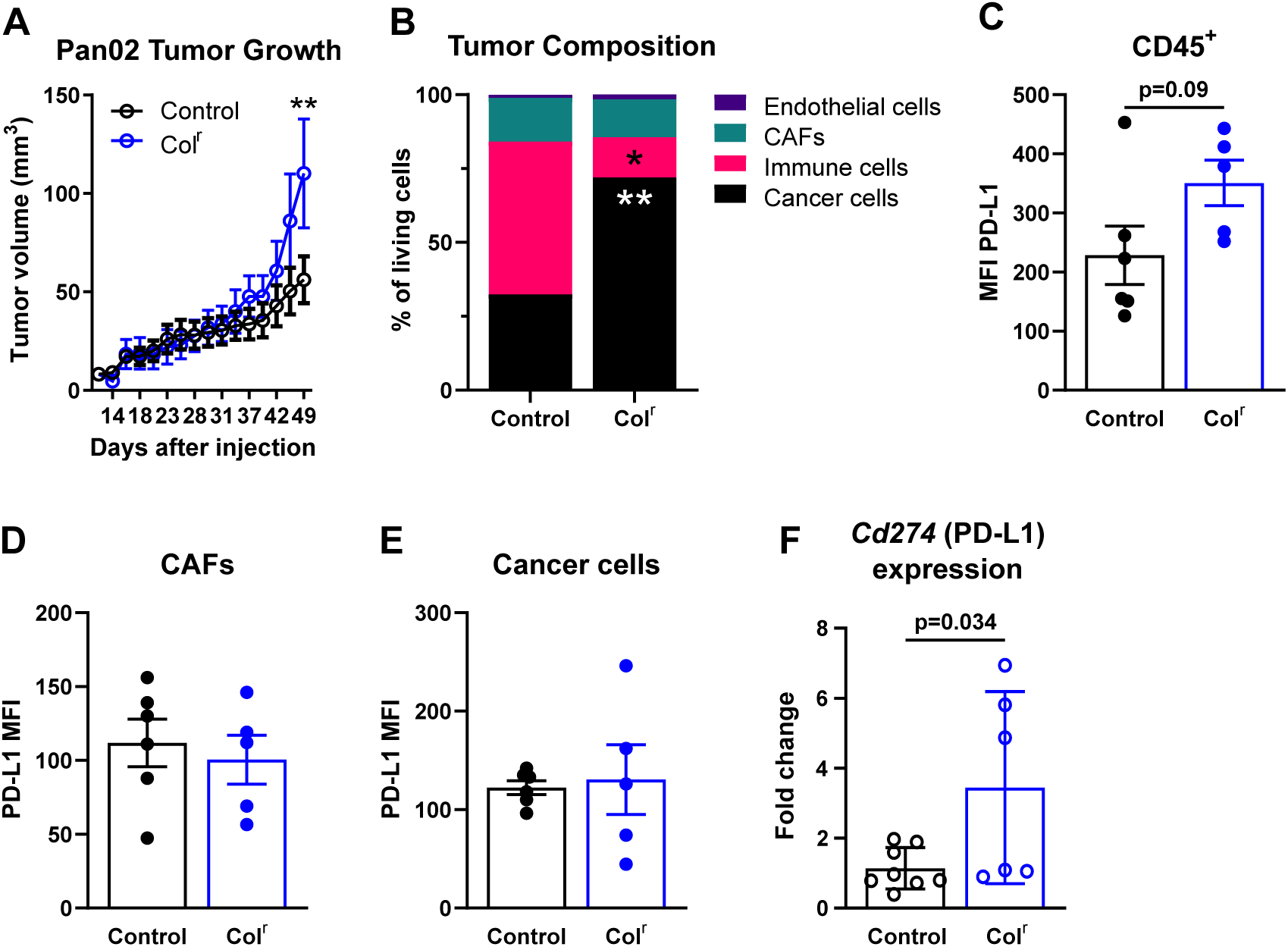
Collagen accumulation increases growth of Pan02 tumors and reduces immune cell infiltration. **(A)** Mean Pan02 tumor growth curves in wildtype littermate control and Col^r^ and mice (n = 13 per group). **(B)** Tumor composition quantified by flow cytometry: cancer cells (CD45^-^FAP^-^CD31^-^), immune cells (CD45^+^), CAFs (CD45^-^FAP^+^PDGFRα^+^; CD45^-^FAP^-^PDGFRα^+^; CD45^-^ FAP^+^PDGFRα^-^), and endothelial cells (CD45^-^CD31^+^) (n = 5-6 per group). **(C-E)** MFI of PD-L1 on immune cells **(C)**, CAFs **(D)** and cancer cells **(E)**. **(F)** *Cd274* gene expression in tumors from control and Col^r^ mice. Statistical significance was assessed using two-way ANOVA with Bonferroni correction (A) and two-tailed Student’s t-test (C-F). Bars represent mean ± SEM (A-E) or SD (F). \**p*<0.05, \*\**p*<0.01.

We proceeded to study the tumor-infiltrating immune cells in more detail using flow cytometry, and interestingly we observed a dramatic decrease in the amount of CD3^+^ T cells (fig. 6A) in the collagen-dense tumors of the Col^r^ mice, and thus in both CD4^+^ and CD8^+^ T cells (fig. 6B-C). Opposite to the observation in Col1a1 KO tumors, the CD4/CD8 ratio was increased in the tumors of Col^r^ mice (fig. 6D). Additionally, we observed fewer CD19^+^ B cells in collagen-dense Col^r^ tumors (fig. 6E) and a trend towards fewer NK cells (fig. 6F). The increased CD4/CD8 ratio and trend of fewer NK cells suggest that increased intratumoral collagen has immunosuppressive effects. Using 3D cell culture assays, we have previously shown that a high collagen density reduces cytotoxic T cell activity ^38^. Using similar 3D cell culture, we examined if collagen density, in addition to reducing NK cell numbers in tumors, could also modulate NK cell activity. However, NK cells did not acquire a dramatically different gene expression profile when cultured in a high-density collagen matrix compared to a low-density collagen matrix (Fig. S3A). In alignment with this, NK cells did not show altered anti-tumor cytotoxic activity upon culture in high- or low-density collagen (Fig. S3B).

**Figure 6.**
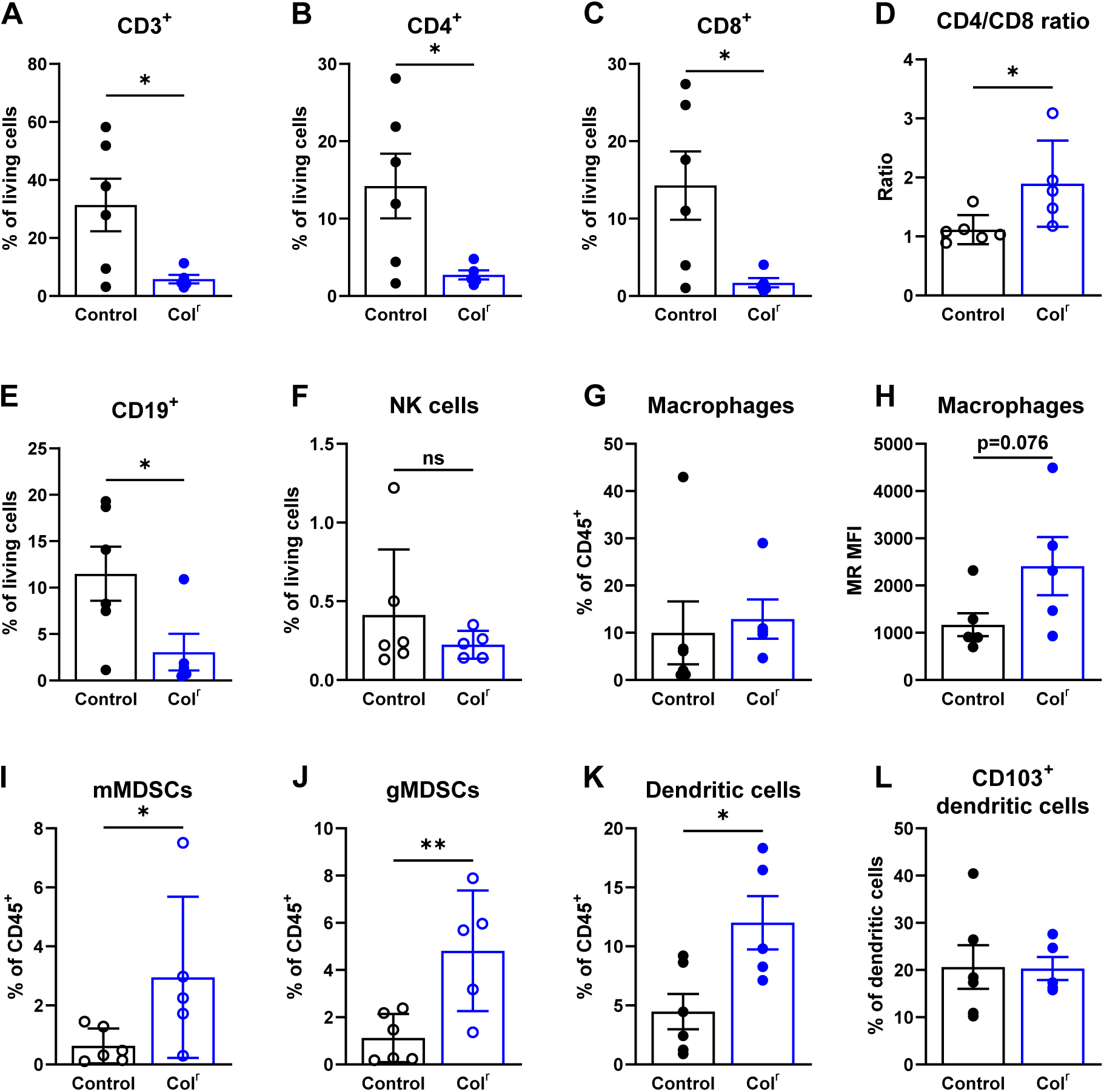
Collagen accumulation decreases T cell infiltration and promotes an immunosuppressive myeloid microenvironment. Pan02 tumor composition in control and Col^r^ mice quantified by flow cytometry: **(A)** T cells (CD3^+^), **(B)** CD4^+^ T cells, **(C)** CD8^+^ T cells, **(D)** CD4/CD8 ratio, **(E)** B cells (CD19^+^), **(F)** NK cells (NK1.1^+^), **(G)** macrophages (CD11b^+^F4/80^+^), **(H)** MFI of MR on macrophages, **(I)** mMDSCs (CD11b^+^F4/80^-^CD11c^-^Ly-6C^+^Ly-6G^-^), **(J)** gMDSCs (CD11b^+^F4/80^-^CD11c^-^Ly-6C^+^Ly-6G^+^), **(K)** dendritic cells (CD11b^+^F4/80^-^CD11c^+^), and **(L)** CD103^+^ dendritic cells. Statistical significance was assessed using the two-tailed Student’s t-test. Bars represent mean ± SEM. \**p*<0.05, \*\**p*<0.01.

When investigating the composition of the myeloid compartment in tumors from Col^r^ mice, the tendencies were largely opposite to observations in Col1a1 KO mice (compare fig. 4 and fig. 6G-L). Macrophage abundance was not affected (fig. 6G) and neither was the expression of MR (fig. 6H), although there was a trend of increased expression in Col^r^ mice, suggesting a slight shift in macrophage polarization towards an immunosuppressive phenotype. The abundance of mMDSCs and gMDSCs were increased in tumors from Col^r^ mice (fig. 6I-J). Dendritic cell numbers were also increased (fig. 6K) although levels of CD103^+^ DCs were not affected (Fig. 6L). In summary, the changes in the myeloid compartment suggest that increased collagen levels induce the formation of a more immunosuppressive TME. The observations in the Col^r^ mice corresponds well with the observations made in the Col1a1 KO mice, which indicate a less immunosuppressive TME in tumors of low collagen content.

To investigate whether a similar correlation between intratumoral collagen levels and immune infiltration exists in human cancer patients, we examined publicly available data from The Cancer Genome Atlas (TCGA). RNA sequencing data from seven cancer types were used to plot the hazard ratios associated with collagen type I expression, defined by a combined COL1A1/COL1A2 signature (fig. 7A). Higher COL1A1/COL1A2 expression was significantly associated with decreased overall survival in skin cutaneous melanoma (SKCM), kidney renal clear cell carcinoma (KIRC), pancreatic adenocarcinoma (PAAD), and bladder urothelial carcinoma (BLCA). Although the associations in lung adenocarcinoma (LUAD), colon adenocarcinoma (COAD), and head and neck squamous cell carcinoma (HNSCC) were not statistically significant, hazard ratios in these cancers showed a trend toward decreased overall survival. In the same TCGA cohorts, we estimated the immune cell composition using CIBERSORTx ^52^ in the top and bottom quartiles of COL1A1/COL1A2 gene expression. Consistent with our findings in the mouse models, tumors with a high collagen type I signature showed a decreased infiltration of CD8 T cells (fig. 7B) and no systematic changes in CD4 T cell infiltration (fig. 7C). An estimated decrease in the abundance of tumor infiltrating activated NK cells was also observed across all seven cancers (fig. 7D), whereas resting NK cells were largely unaffected (fig. 7E). In contrast, the abundance of TAMs was increased in tumors with a high COL1A1/COL1A2 expression in all seven cohorts (fig. 7F). These data support a central role for intratumoral collagen in controlling immune cell infiltration and activity and thereby promoting an immunosuppressive tumor microenvironment.

**Figure 7.**
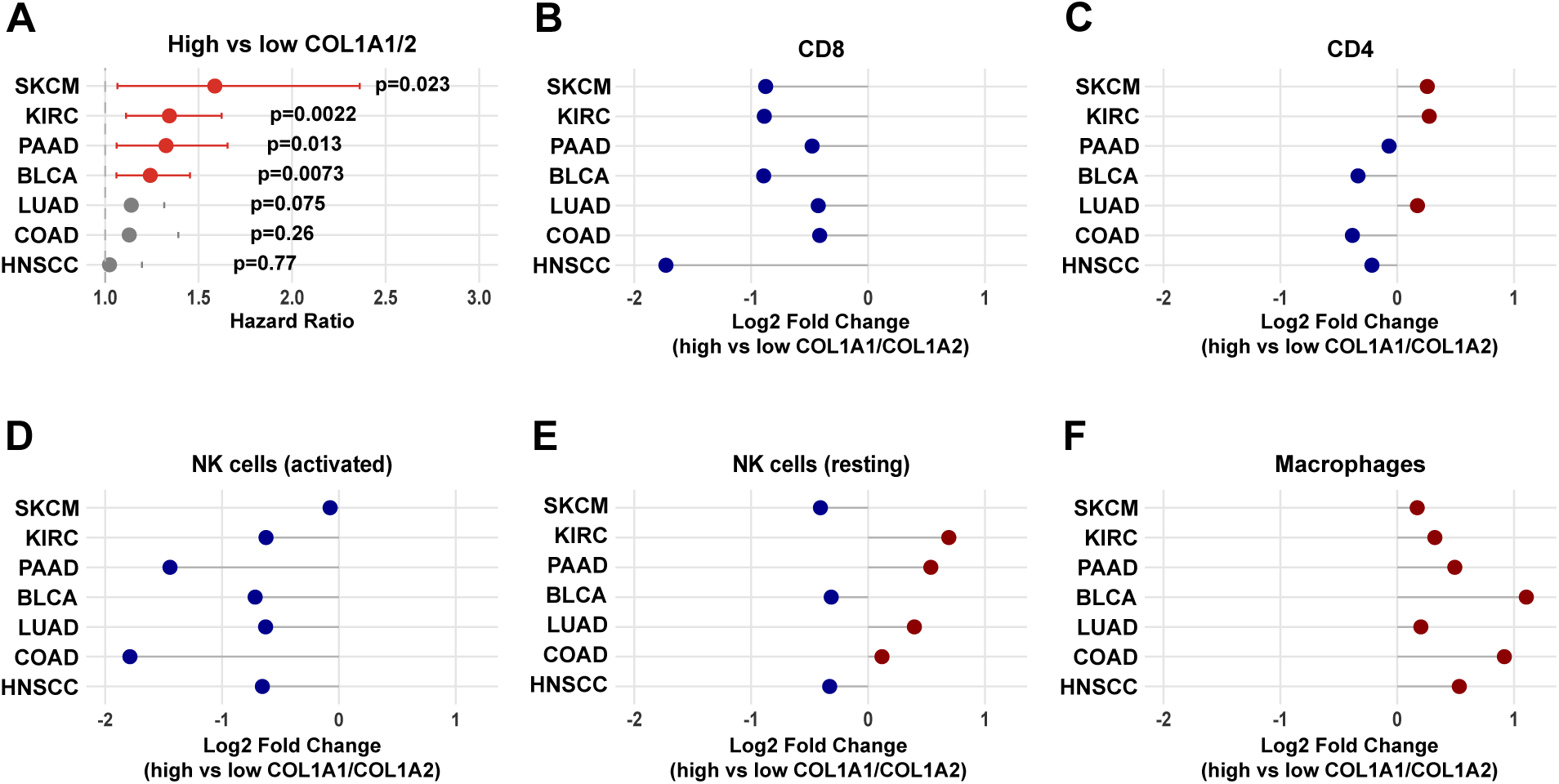
Collagen type I expression levels are associated with altered immune infiltration in human cancers. **(A)** Hazard ratios with 95% confidence intervals for the COL1A1/COL1A2 combined expression signature, obtained from Cox proportional hazards models restricted to primary tumor samples from TCGA covereing the following cancer types: SKCM, KIRC, PAAD, BLCA, LUAD, COAD and HNSCC. P-values correspond to the coefficient significance reported by the Cox model. **(B-F)** Plots showing differences in fraction change of CD8+ T cells **(B)**, CD4+ T cells **(C)**, activated NK cells **(D)**, resting NK cells **(E)**, and Macrophages **(F)** between the top and bottom quartiles of the COL1A1/COL1A2 combined expression signature in primary tumor samples from TCGA cohorts. Differences are displayed as log₂ fold change (High/Low COL1A1/COL1A2), with positive values indicating higher fraction in the top quartile.

## Discussion

Increased content and alignment of collagen type I in tumors have been suggested as markers for poor prognosis in many types of cancer ^21–25^. However, the underlying mechanisms behind these correlations are still not uncovered. In this study, we employed an inducible systemic Col1a1 KO mouse model to directly investigate the impact of collagen on tumor progression and immune cell infiltration. In order to effectively reduce collagen type I levels in tumors, we generated β-actin-cre/ER;Col1a1^flox/flox^ mice in which tamoxifen treatment induces global *Col1a1* gene inactivation. We hypothesized that the lack of Col1 neosynthesis could be tolerated in adult mice due to the slow turnover of pre-existing collagen fibers. In our study we found that the mouse model worked as intended since tamoxifen treatment led to an approximately 80% reduction in Col1a1 transcript levels, which in turn caused a marked decrease in intratumoral collagen levels. The tamoxifen-treated mice eventually developed signs of reduced joint flexibility, and within four weeks of tamoxifen exposure, all mice had to be euthanized due to impaired mobility and progressive weight loss. However, within this time period, we could demonstrate that reduced collagen levels led to impeded growth of subcutaneous Pan02 tumors. In further support of a pro-tumorigenic role for collagen, we also observed that increased collagen content in transgenic Col^r^ mice led to accelerated tumor growth.

Tumors can be classified according to their degree of immune cell infiltration as “inflamed”, “immune excluded”, or “immune desert” ^53^. In immune-excluded tumors, immune cells are typically found in the stroma surrounding the tumor cell nests. Studies investigating effects of collagen density and stiffness on immune cell migration have shown that T cells migrate more efficiently in a stroma with a loose network of collagen compared to a dense collagen network, and that reducing stiffness through inhibition of collagen crosslinking improves T cell migration ^35,54^. In line with these findings, we observed a dramatic decrease in the number of tumor-infiltrating immune cells in the collagen-dense tumors of Col^r^ mice, with T cells being most notably affected. Tumor-infiltration of CD8^+^ T cells seemed to be more impaired by collagen than CD4^+^ T cells, leading to an increased CD4/CD8 ratio in Col^r^ mice and a decreased CD4/CD8 ratio in Col1a1 KO mice. In addition, we found a significant increase in the abundance of tumor-infiltrating NK cells in Col1a1 KO mice.

In earlier work, we demonstrated that conditioned medium from macrophages cultured in high-density collagen gels impaired migration of T cells, with CD8+ T cells being particularly affected ^37^. This reduced migration of CD8^+^ T cells in high collagen densities could explain our observed decrease in CD4/CD8 ratio in Col1a1 KO mice and increase in CD4/CD8 ratio in Col^r^ mice. A similar effect of collagen has also been shown in a murine *in vivo* model of wound repair, where collagen scaffolds promoted a higher CD4/CD8 ratio compared to saline-treated controls ^55^. Using 3D culture models, we previously demonstrated that a high collagen density also impairs T cell proliferation ^38^. It remains to be determined whether reduced proliferation also contributes to the decreased abundance of tumor-infiltrating lymphocytes observed in the current study. Interestingly, a study by Perez-Penco *et al.* showed that modulation of the TME of PAN02 tumors using a TGFβ-derived peptide vaccine led to reduced collagen production accompanied by increased infiltration of CD8 T cells, in line with our findings in the current study ^56^.

We also previously showed that a high collagen density reduces the cytotoxic capacity of tumor-infiltrating T cells against autologous cancer cells in 3D culture models, highlighting that collagen can support immune evasion by cancer cells ^28^. Here, we sorted T cells from tumors from Col1a1 KO mice and wildtype littermates and analyzed the gene expression of a limited number of markers of T cell activity and exhaustion. This result suggests that the pro-tumorigenic effects of collagen include modulation of the cytotoxic activity of infiltrating T cells although more studies are required to fully elucidate this. The observed collagen-dependent changes in the numbers of tumor-infiltrating NK cells prompted us to examine in a 3D cell culture model if a high collagen density could lead to reduced NK cell activity. However, NK cells did not seem to respond to the surrounding collagen density in a similar manner as previously observed for T cell ^38^.

Myeloid cell populations within the TME were also affected by collagen density. Specifically, we observed a reduction in the number of mMDSCs in low-collagen tumors of Col1a1 KO mice and a trend towards more mMDSCs in collagen-dense tumors of Col^r^ mice. There was also a marked increase in gMDSCs in collagen-dense tumors. In light of these results, it is noteworthy that, García-Mendoza *et al.* reported that the cytokine GM-CSF was upregulated in collagen-dense mammary tumors of Colr mice. Given the known ability of this cytokine to promote the infiltration of myeloid cells into tumors ^27^, GM-CSF may also contribute to the collagen-dependent effects on the myeloid cell compartment observed in our pancreatic tumor models.

Recently, another conditional Col1 KO mouse model was used by Chen *et al.* to study collagen-mediated effects on tumor progression ^41^. In that study the *Col1a1* gene was inactivated specifically in αSMA myofibroblasts, which allowed them to perform long-running experiments at the expense of a lower knockout efficiency at around 50%. Surprisingly, Col1 deletion in this study resulted in decreased overall survival in a genetically induced mouse model of pancreatic cancer. The underlying mechanism was proposed to involve enhanced recruitment of immunosuppressive myeloid cells in the tumors of conditional col1 KO mice. Another study investigated the role of collagen for liver metastasis growth and similarly observed increased tumor progression upon reduction of col1 levels ^40^. This reduction was achieved through hepatic stellate cell–specific or inducible liver-specific deletion of Col1a1. The anti-tumorigenic effect of collagen in that context was suggested to involved encapsulation of the tumor, a process reminiscent of bacterial abscess formation ^40^. These results stand in stark contrast to the observations of this study. While differences in KO strategies and tumor models may account for some of the discrepancies, the divergent effects reported by Chen *et al.* are particularly notable. In contrast to their findings, we observed decreased pancreatic tumor growth in Col1a1 KO mice as well as a significant decrease in mMDSCs and reduced PD-L1 expression, indicating a less immunosuppressive TME.

Interestingly, studies by Tian *et al.* have demonstrated that cancer cell-derived collagen, but not stroma-derived collagen, suppress PDAC growth ^57,58^. Such a distinction between stroma-derived and cancer cell-derived collagen could be one of the underlying reasons for the seemingly contradictory results. Taken together, these studies emphasize the complexity of collagen’s role in tumor progression and highlight the importance of considering the cellular source of collagen and the tumor context.

In conclusion, by using both a conditional Col1a1 KO mouse model and a collagen-accumulating mouse model, we demonstrate that collagen density affects tumor growth and the level of tumor-infiltrating immune cells in a murine model of pancreatic cancer. The controversy regarding the tumor-suppressive or -supportive role of collagen in tumor progression highlights the need for further research to dissect the precise and context-dependent functions of collagen in cancer progression.

## Experimental Procedures

### Cell culture

The murine pancreatic tumor cell line Pan02 was obtained from the NCI Repository and cultured in cell culture treated flasks (Corning Incorporated) in DMEM + GlutaMAXTM-I (Gibco) supplemented with 10% fetal bovine serum (FBS) (Life Technologies) and 1% penicillin and streptomycin (P/S) (Gibco). Prior to injection into mice, cells were detached by incubation with trypsin-EDTA (Gibco) for 5 min at 37°C, washed with phosphate buffered saline (PBS) (Gibco), centrifuged at 300xg for 5 min and resuspended in unsupplemented DMEM.

### Animal experiments

All animal experiments were performed according to institutional guidelines and under licenses issues by the Animal Experiments Inspectorate in Denmark. Col1a1^flox/flox^ mice have previously been described ^42^. They were interbred with β-actin-cre/ER mice (The Jackson Laboratory, strain 004682) to generate mice that were either positive for β-actin-cre/ER (Col1a1 KO) or negative (control) and all were homozygous for Col1a1^flox/flox^. Col1a1 deletion was induced by administration of 200µL 20mg/mL tamoxifen (Sigma-Aldrich) by oral gavage for 5 consecutive days followed by three days per week until the end of the experiment. Tamoxifen was dissolved in corn oil (Sigma-Aldrich) by vigorous vortexing and incubation at 37°C overnight.

Col1a1^tm1jae/+^ mice (The Jackson Laboratory, strain 002495) were interbred with wildtype C57BL/6 mice (Taconic) to generate heterozygous Col1a1^tm1jae/+^ (Col^r^) mice and wildtype littermate control mice.

All mice were genotyped by PCR on DNA extracted from ear clippings. Col1a1 KO mice and wildtype littermates were genotyped for the presence of the β-actin-cre transgene (primers Cre_fw, 5′-GCGGTCTGGCAGTAAAAACTATC-3′ and Cre_rv, 5′-GTGAAACAGCATTGCTGTCACTT-3′). Successful reaction was confirmed using primers Int_Con_fw, 5′-CAAATGTTGCTTGTCTGGTG-3′ and Int_Con_rv, 5′-GTCAGTCGAGTGCACAGTTT-3′. For analysis of the floxed *Col1a1*, the following primers were used: Col_flox_fw: 5′-TGGTACAGCACTTTACAGCGCACA-3′ and Col_flox_rv: 5′-TTACTCGGCCTGGGTCACTTCTTT-3′. The reaction for genotyping of Col1a1 mice (Cre and control) was as follows: 95°C for 5 min, 35 cycles of 94°C for 45 sec, 60°C for 30 sec and 72°C for 60 sec, followed by 72°C for 10 min. The reaction for floxed *Col1a1* was as follows: 95°C for 5 min, 33 cycles of 94°C for 45 sec, 55°C for 45 sec and 72°C for 60 sec, followed by 72°C for 10 min. Col^r^ mice were genotyped for the wildtype *Col1a1* allele (Col1a1wt-1=5′-TGGACAACGTGGTGTGGTC-3′ and Col1a1-wt-2=5′-TTGAACTCAGGAATTTACCTGC-3′) and the mutated allele (Col1a1mut-1= 5′-TGGACAACGTGGTGCCGCG-3′ and Col1a1wt-2) using the following reaction: 95°C for 5 min, 35 cycles of 95°C for 30 sec, 60°C for 30 sec and 72°C for 60 sec, followed by 72°C for 10 min.

For tumor studies, 0.5*10^6^ Pan02 cells were injected in 100µL of DMEM subcutaneously on one flank. Tumors were measured three times per week using calipers and tumor volumes were calculated using the formula: (length * width^2^) * 0.5. Mice treated with ICI were injected intraperitoneally three times per week from when tumors became palpable and until endpoint. They were injected with a mixture of anti-PD-L1 mAb (200µg/mouse, clone 10F.9G2, BioXcell) and anti-CTLA4 mAb (200 μg/mouse; clone 9D9, BioXcell).

### Multicolor flow cytometry

After harvest, tumors were finely diced and dissociated by incubation in digestion buffer (75μg/mL DNase I (Worthington) and (2.1 mg/mL bacterial collagenase type I (Corning) in RPMI (Gibco by Life Technologies) supplemented with 1% Penicillin-Streptomycin) at 37°C with rotation for 1 hour. Then, the suspension was homogenized by thorough pipetting before being filtered through 70 μm cell strainers (Corning) and centrifuged at 300xg for 5 min. To lyse erythrocytes, the pellet was resuspended in RBC lysis buffer (Sigma-Aldrich), pipetted gently up and down for 30 seconds and left still for 2 min before RPMI with 10% FBS and 1% P/S was added to dilute the RBC lysis buffer. The cells were centrifuged at 300xg for 5 min and resuspended in staining buffer (5mM EDTA, 0.5% bovine serum albumin in PBS (Lonza)) containing FcR-blocking reagent (Miltenyi Biotec) according to manufacturer’s instructions before staining. All cells or purified CD45^+^ cells were stained with an antibody cocktail containing antibodies for either a general (all cells), myeloid-, or lymphoid panel by incubation for 20 min in the dark at 4°C (see table S1 for details on antibody panels). The cells were then washed in staining buffer and centrifuged at 300xg for 5 min. The supernatant was discarded, and the cells resuspended in 250μL staining buffer. Before resuspension in 250μL staining buffer, samples stained with the general panel underwent a secondary staining with streptavidin-APC (Biolegend) for 10 min at 4°C in the dark. The samples were analyzed using a FACSCantoII (BD Biosciences) or a Novocyte Quanteon (Agilent) and data was analyzed with FlowJo Software. Fluorescence minus one (FMO) and isotype controls were included were appropriate. All panels underwent a compensation procedure prior to analysis. All antibodies were from Biolegend unless otherwise stated: Brilliant Violet 421™ anti-mouse CD274 (B7-H1, PD-L1), FITC anti-mouse CD31, PE/Cy7 anti-mouse CD45, PE anti-mouse CD140a (PDGFRa), FAP Biotinylated Antibody (R&D Systems), Brilliant Violet 421™ anti-mouse CD103, FITC anti-mouse CD11c, PE anti-mouse CD206 (MR), PerCP/Cy5.5 anti-mouse Ly-6C, PE/Cy7 anti-mouse/human CD11b, APC anti-mouse F4/80, APC/Cy7 anti-mouse Ly-6G, Brilliant Violet 421™ anti-mouse CD4, FITC anti-mouse CD3, PE anti-mouse CD279 (PD-1), PerCP/Cy5.5 anti-mouse CD25, PE-Cy7 anti-mouse CD19, APC anti-mouse CD8a, and APC-Cy7 anti-mouse NK1.1. Dead cells were excluded using Zombie Aqua Dye (Biolegend).

### CD45+ cell enrichment

Following FcR-blocking as described above, cell suspensions were enriched for the CD45^+^ fraction using anti-CD45 MicroBeads (Miltenyi Biotec) according to the manufacturer’s instructions.

### Fluorescence-activated cell sorting (FACS)

Following CD45 enrichment, CD4^+^ and CD8^+^ T cells were sorted. The CD45^+^ fraction was stained with Zombie Aqua™ Fixable Viability Kit, FITC anti-mouse CD3, Brilliant Violet 421™ anti-mouse CD4, and APC anti-mouse CD8a as described above. Cells were sorted into RPMI plus 50% FBS using a FACSMelody (BD Biosciences). RNA was isolated immediately after sorting.

### RNA isolation

After sorting, the isolated cells were centrifuged for 5 min at 300xg and lysed by resuspension in 350 μL RLT buffer supplemented with β-mercaptoethanol. RNA was isolated using the RNeasy Mini Kit (Qiagen) according to the manufacturer’s instructions. The optional treatment with DNase was included. Quantity of the purified RNA was determined using an Agilent 2100 BioAnalyzer (Agilent Genomics).

### Quantitative real-time PCR

cDNA was synthesized from RNA using the iScript cDNA Synthesis Kit (Bio-Rad Laboratories) according to the manufacturer’s instructions. Controls without reverse transcriptase or without template were included. The quantitative real-time PCR (qRT-PCR) was done using the Brilliant III Ultra-Fast SYBR Green QPCR Master Mix (Agilent Technologies) according to the manufacturer’s instructions. The program used was as follows: 3 min at 95°C, 40 cycles of 5 seconds at 95°C, 40 cycles of 20 seconds at 60°C, and lastly a melting curve analysis of 65-95°C with 0.5°C increment, 5 seconds per step. qRT-PCR was done using an AriaMX Real-Time PCR System (G8830A). Everything was run in quadruplicates and normalized to the internal reference gene, *Actb*. Relative fold changes were calculated using the comparative cycle threshold (ΔΔCT) method. Primers were designed using the Primer-BLAST tool (National Center for Biotechnology Information, National Institutes of Health). Primer sequences are listed in table S2.

### Histological staining

Tumors were fixed overnight in formalin at room temperature, then transferred to 70% ethanol and stored at 4°C until paraffin embedding. Samples were embedded in paraffin before being cut into 4μm sections. Sections were stained with picrosirius red (Ampliqon) according to manufacturer’s instructions or Masson’s trichrome according to manufacturer’s instructions. Images of PSR stained slides were acquired using a polarization filter. Positive areas were quantified using Qupath software (ver. 0.2.3) using the Qupath Pixel Classifier ^59^.

### Competitive ELISA

Blood was collected from the mice immediately after euthanasia and incubated at room temperature for 30 min. Samples were centrifuged for 10 min at 1500xg and the serum was carefully transferred to clean tubes and frozen at −80°C. The MMP-generated Col1 degradation product, C1M, and PRO-C1, which is an internal epitope from the N-terminal propeptide of Col1, were measured using competitive ELISAs developed by Nordic Bioscience ^60,61^. Briefly, 96-well plates precoated with streptavidin were coated with biotinylated peptides specific for C1M and PRO-C1, respectively, and incubated at 20°C for 30 min. Standard peptide or serum samples were added followed by addition of appropriate peroxidase-conjugated monoclonal antibodies and incubated at 2-8°C for 20 hours for C1M and for 3 hours for PRO-C1. Then, tetramethylbenzinidine (Kem-En-Tec Diagnostics) was added to the plate and incubated at 20°C for 15 min. Next, the reaction was stopped by addition of 1% sulfuric acid and absorbance was measured at 450 nm with 650 nm as reference. Analysis was performed using the Softmax Pro v. 6.3 software. Values were assigned the lower level of detection if they were below.

### TCGA data analysis

RNA sequencing data and corresponding clinical metadata from skin cutaneous melanoma (SKCM), colon adenocarcinoma (COAD), kidney renal clear cell carcinoma (KIRC), pancreatic adenocarcinoma (PAAD), bladder urothelial carcinoma (BLCA), head and neck squamous cell carcinoma (HNSCC), and lung adenocarcinoma (LUAD) were obtained from The Cancer Genome Atlas (TCGA) via the UCSC Xena Browser. Only primary tumor samples (sample barcodes ending in 01A) were included. Clinical follow-up time was calculated as the number of days from diagnosis to death or last follow-up. Samples lacking a recorded date of death were assigned the reported time to last follow-up, and follow-up times were right-censored at 1,825 days (5 years).

Gene expression values for COL1A1 and COL1A2 were extracted and combined into a single signature score calculated as the mean of the two genes. Cox proportional hazards models were fitted using the COL1A1/COL1A2 signature as a continuous variable to estimate hazard ratios (HRs) and 95% confidence intervals for each TCGA cohort. HRs were visualized individually and compiled into a combined forest plot summarizing the association between COL1A1/COL1A2 expression and overall survival across all seven cancer types. All analyses were performed in R (version 4.5.0). Immune cell composition was estimated using CIBERSORTx ^52^ with the LM22 signature matrix on primary tumor samples (barcodes ending in 01A) from the same TCGA cohorts used for hazard ratio analysis. For each sample, 100 permutations were performed to assess significance, and only samples with p < 0.05 were retained for downstream analysis. The combined ECM signature score (mean of COL1A1 and COL1A2 expression) was calculated for each sample, and samples were stratified into top and bottom quartiles. Immune cell fractions were compared between these groups. CD4 T cells were combined from Memory Activated, Memory Resting, Naive, Follicular Helper, and Regulatory T cell fractions. Macrophage groups were combined from Macrophages M0, Macrophages M1, and Macrophages M2 fractions. CD8 and NK cell populations were plotted as individual samples.

Differences in immune cell fractions between top and bottom ECM signature quartiles were visualized as log₂ fold change using lollipop plots. Positive values indicate a higher fraction in the top quartile. All analyses and plotting were completed in R (version 4.5.0).

### Statistical analyses

All statistical tests were performed using GraphPad Prism 8. For comparison of two groups, unpaired two-tailed Student’s t-test was used. For analysis of TCGA data, statistical significance between quartiles was determined using the Wilcoxon rank-sum test. To determine statistical significance for tumor growth with multiple comparison between multiple timepoints, two-way ANOVA with Bonferroni correction was used.

## Author contributions

Conceptualization, M-L.T. and D.H.M.; Methodology, M-L.T., L.H.E., N.W., K.K., M.H.A., L.G. and D.H.M.; Investigation, M-L.T., A.Z.J., K.J.B, M.C., A.M.A.R, C.J., H.J.J., H.L., N.K.C, D.E.K., K.Y.F., and D.H.M; Writing – Original Draft, M-L.T. and D.H.M.; Writing – Review & Editing, , M-L.T., A.Z.J., K.J.B, M.C., A.M.A.R, C.J., H.J.J., H.L., N.K.C, D.E.K., K.Y.F., L.H.E., N.W., K.K., M.H.A., L.G. and D.H.M.

## Acknowledgement

This work was supported by the Lundbeck Foundation (R307-2018-3326) (to D.H.M.), Danish Cancer Society (R231-A14035 (to D.H.M.), and Department of Oncology, Copenhagen University Hospital – Herlev & Gentofte.

## References

1 Pickup MW, Mouw JK, Weaver VM. The extracellular matrix modulates the hallmarks of cancer. EMBO Rep 2014; 15: 1243–53.

2 Fang M, Yuan J, Peng C, Li Y. Collagen as a double-edged sword in tumor progression. Tumour Biol 2014; 35: 2871–2882.

3 Levental KR, Yu H, Kass L, Lakins JN, Egeblad M, Erler JT et al. Matrix Crosslinking Forces Tumor Progression by Enhancing Integrin Signaling. Cell 2009; 139: 891–906.

4 Schedin P, Keely PJ. Mammary gland ECM remodeling, stiffness, and mechanosignaling in normal development and tumor progression. Cold Spring Harb Perspect Biol 2011; 3: a003228.

5 Ray A, Callaway MK, Rodríguez-Merced NJ, Crampton AL, Carlson M, Emme KB et al. Stromal architecture directs early dissemination in pancreatic ductal adenocarcinoma. JCI Insight 2022; 7. doi:10.1172/JCI.INSIGHT.150330.

6 Rømer AMA, Thorseth ML, Madsen DH. Immune Modulatory Properties of Collagen in Cancer. Front Immunol. 2021; 12. doi:10.3389/fimmu.2021.791453.

7 Naba A, Clauser KR, Lamar JM, Carr SA, Hynes RO. Extracellular matrix signatures of human mammary carcinoma identify novel metastasis promoters. Elife 2014; 3: e01308.

8 Gelse K, Pöschl E, Aigner T. Collagens - Structure, function, and biosynthesis. Adv Drug Deliv Rev 2003; 55: 1531–1546.

9 Kadler KE, Baldock C, Bella J, Boot-Handford RP. Collagens at a glance. J Cell Sci 2007; 120: 1955–1958.

10 Shoulders MD, Raines RT. Collagen structure and stability. Annu Rev Biochem 2009; 78: 929–958.

11 Song F, Wisithphrom K, Zhou J, Windsor LJ. Matrix metalloproteinase dependent and independent collagen degradation. Front Biosci 2006; 11: 3100–3120.

12 Highberger JH, Corbett C, Gross J. Isolation and characterization of a peptide containing the site of cleavage of the chick skin collagen alpha 1[I] chain by animal collagenases. Biochem Biophys Res Commun 1979; 89: 202–8.

13 Madsen DH, Bugge TH. The source of matrix-degrading enzymes in human cancer: Problems of research reproducibility and possible solutions. J Cell Biol 2015; 209: 195–8.

14 Madsen DH, Engelholm LH, Ingvarsen S, Hillig T, Wagenaar-Miller RA, Kjøller L et al. Extracellular collagenases and the endocytic receptor, urokinase plasminogen activator receptor-associated protein/endo180, cooperate in fibroblast-mediated collagen degradation. Journal of Biological Chemistry 2007; 282: 27037–27045.

15 McKleroy W, Lee TH, Atabai K. Always cleave up your mess: targeting collagen degradation to treat tissue fibrosis. Am J Physiol Lung Cell Mol Physiol 2013; 304. doi:10.1152/AJPLUNG.00418.2012.

16 Murphy G, Reynolds JJ, Bretz U, Baggiolini M. Partial purification of collagenase and gelatinase from human polymorphonuclear leucocytes. Analysis of their actions on soluble and insoluble collagens. Biochem J 1982; 203: 209–221.

17 Madsen DH, Ingvarsen S, Jürgensen HJ, Melander MC, Kjøller L, Moyer A et al. The non-phagocytic route of collagen uptake: A distinct degradation pathway. Journal of Biological Chemistry 2011; 286: 26996–27010.

18 Madsen DH, Jürgensen HJ, Siersbæk MS, Kuczek DE, Grey Cloud L, Liu S et al. Tumor-Associated Macrophages Derived from Circulating Inflammatory Monocytes Degrade Collagen through Cellular Uptake. Cell Rep 2017; 21: 3662–3671.

19 Jürgensen HJ, van Putten S, Nørregaard KS, Bugge TH, Engelholm LH, Behrendt N et al. Cellular uptake of collagens and implications for immune cell regulation in disease. Cell Mol Life Sci 2020; 77: 3161–3176.

20 Thorseth ML, Carretta M, Jensen C, Mølgaard K, Jürgensen HJ, Engelholm LH et al. Uncovering mediators of collagen degradation in the tumor microenvironment. Matrix Biol Plus 2022; 13. doi:10.1016/j.mbplus.2022.100101.

21 Esbona K, Yi Y, Saha S, Yu M, Van Doorn RR, Conklin MW et al. The Presence of Cyclooxygenase 2, Tumor-Associated Macrophages, and Collagen Alignment as Prognostic Markers for Invasive Breast Carcinoma Patients. American Journal of Pathology 2018; 188: 559–573.

22 Drifka CR, Loeffler AG, Mathewson K, Keikhosravi A, Eickhoff JC, Liu Y et al. Highly aligned stromal collagen is a negative prognostic factor following pancreatic ductal adenocarcinoma resection. Oncotarget 2016; 7: 76197–76213.

23 Li H-X, Zheng J-H, Fan H-X, Li H-P, Gao Z-X, Chen D. Expression of αvβ6 integrin and collagen fibre in oral squamous cell carcinoma: association with clinical outcomes and prognostic implications. Journal of Oral Pathology & Medicine 2013; 42: 547–556.

24 Ohno S, Tachibana M, Fujii T, Ueda S, Kubota H, Nagasue N. Role of stromal collagen in immunomodulation and prognosis of advanced gastric carcinoma. Int J Cancer 2002; 97: 770–774.

25 Conklin MW, Eickhoff JC, Riching KM, Pehlke CA, Eliceiri KW, Provenzano PP et al. Aligned collagen is a prognostic signature for survival in human breast carcinoma. American Journal of Pathology 2011; 178: 1221–1232.

26 Provenzano PP, Inman DR, Eliceiri KW, Knittel JG, Yan L, Rueden CT et al. Collagen density promotes mammary tumor initiation and progression. BMC Med 2008; 6: 11.

27 García-Mendoza MG, Inman DR, Ponik SM, Jeffery JJ, Sheerar DS, Van Doorn RR et al. Neutrophils drive accelerated tumor progression in the collagen-dense mammary tumor microenvironment. Breast Cancer Res 2016; 18. doi:10.1186/S13058-016-0703-7.

28 Barcus CE, O’Leary KA, Brockman JL, Rugowski DE, Liu Y, Garcia N et al. Elevated collagen-I augments tumor progressive signals, intravasation and metastasis of prolactin-induced estrogen receptor alpha positive mammary tumor cells. Breast Cancer Res 2017; 19. doi:10.1186/S13058-017-0801-1.

29 Esbona K, Inman D, Saha S, Jeffery J, Schedin P, Wilke L et al. COX-2 modulates mammary tumor progression in response to collagen density. Breast Cancer Res 2016; 18. doi:10.1186/S13058-016-0695-3.

30 Liu X, Wu H, Byrne M, Jeffrey J, Krane S, Jaenisch R. A targeted mutation at the known collagenase cleavage site in mouse type I collagen impairs tissue remodeling. J Cell Biol 1995; 130: 227–237.

31 Paszek MJ, Zahir N, Johnson KR, Lakins JN, Rozenberg GI, Gefen A et al. Tensional homeostasis and the malignant phenotype. Cancer Cell 2005; 8: 241–254.

32 Mammoto A, Connor KM, Mammoto T, Yung CW, Huh D, Aderman CM et al. A mechanosensitive transcriptional mechanism that controls angiogenesis. Nature 2009; 457: 1103–1108.

33 Gordon-Weeks A, Yuzhalin AE. Cancer Extracellular Matrix Proteins Regulate Tumour Immunity. Cancers (Basel) 2020; 12: 1–25.

34 Peranzoni E, Rivas-Caicedo A, Bougherara H, Salmon H, Donnadieu E. Positive and negative influence of the matrix architecture on antitumor immune surveillance. Cell Mol Life Sci 2013; 70: 4431–48.

35 Salmon H, Franciszkiewicz K, Damotte D, Dieu-Nosjean M-C, Validire P, Trautmann A et al. Matrix architecture defines the preferential localization and migration of T cells into the stroma of human lung tumors. J Clin Invest 2012; 122: 899–910.

36 Hartmann N, Giese NA, Giese T, Poschke I, Offringa R, Werner J et al. Prevailing Role of Contact Guidance in Intrastromal T-cell Trapping in Human Pancreatic Cancer. Clinical Cancer Research 2014; 20: 3422–3433.

37 Larsen AMH, Kuczek DE, Kalvisa A, Siersbæk MS, Thorseth M-L, Johansen AZ et al. Collagen Density Modulates the Immunosuppressive Functions of Macrophages. The Journal of Immunology 2020; 205: 1461–1472.

38 Kuczek DE, Larsen AMH, Thorseth M-L, Carretta M, Kalvisa A, Siersbæk MS et al. Collagen density regulates the activity of tumor-infiltrating T cells. J Immunother Cancer 2019; 7: 68.

39 Callaway MK, Noonan BJ, Schwertfeger KL, Provenzano PP. Extracellular matrix architecture promotes immunosuppressive microenvironments in pancreatic cancer. Matrix Biol 2025. doi:10.1016/J.MATBIO.2025.09.004.

40 Bhattacharjee S, Hamberger F, Ravichandra A, Miller M, Nair A, Affo S et al. Tumor restriction by type I collagen opposes tumor-promoting effects of cancer-associated fibroblasts. J Clin Invest 2021; 131. doi:10.1172/JCI146987.

41 Chen Y, Kim J, Yang S, Wang H, Wu CJ, Sugimoto H et al. Type I collagen deletion in αSMA+ myofibroblasts augments immune suppression and accelerates progression of pancreatic cancer. Cancer Cell 2021; 39: 548–565.e6.

42 Yang J, Wheeler SE, Velikoff M, Kleaveland KR, Lafemina MJ, Frank JA et al. Activated alveolar epithelial cells initiate fibrosis through secretion of mesenchymal proteins. American Journal of Pathology 2013; 183: 1559–1570.

43 Carretta M, Thorseth ML, Schina A, Agardy DA, Johansen AZ, Baker KJ et al. Dissecting tumor microenvironment heterogeneity in syngeneic mouse models: insights on cancer-associated fibroblast phenotypes shaped by infiltrating T cells. Front Immunol 2023; 14. doi:10.3389/FIMMU.2023.1320614/PDF.

44 Pentcheva-Hoang T, Corse E, Allison JP. Negative regulators of T-cell activation: potential targets for therapeutic intervention in cancer, autoimmune disease, and persistent infections. Immunol Rev 2009; 229: 67–87.

45 Bougherara H, Mansuet-Lupo A, Alifano M, Ngô C, Damotte D, Le Frère-Belda MA et al. Real-Time Imaging of Resident T Cells in Human Lung and Ovarian Carcinomas Reveals How Different Tumor Microenvironments Control T Lymphocyte Migration. Front Immunol 2015; 6. doi:10.3389/FIMMU.2015.00500.

46 Peranzoni E, Ingangi V, Masetto E, Pinton L, Marigo I. Myeloid Cells as Clinical Biomarkers for Immune Checkpoint Blockade. Front Immunol 2020; 11. doi:10.3389/FIMMU.2020.01590.

47 Broz ML, Binnewies M, Boldajipour B, Nelson AE, Pollack JL, Erle DJ et al. Dissecting the Tumor Myeloid Compartment Reveals Rare Activating Antigen-Presenting Cells Critical for T Cell Immunity. Cancer Cell 2014; 26: 938.

48 Salmon H, Idoyaga J, Rahman A, Leboeuf M, Remark R, Jordan S et al. Expansion and Activation of CD103+ Dendritic Cell Progenitors at the Tumor Site Enhances Tumor Responses to Therapeutic PD-L1 and BRAF Inhibition. Immunity 2016; 44: 924–938.

49 Bai R, Lv Z, Xu D, Cui J. Predictive biomarkers for cancer immunotherapy with immune checkpoint inhibitors. Biomark Res 2020; 8. doi:10.1186/S40364-020-00209-0.

50 Luheshi NM, Coates-Ulrichsen J, Harper J, Mullins S, Sulikowski MG, Martin P et al. Transformation of the tumour microenvironment by a CD40 agonist antibody correlates with improved responses to PD-L1 blockade in a mouse orthotopic pancreatic tumour model. Oncotarget 2016; 7: 18508–18520.

51 Marshall LA, Marubayashi S, Jorapur A, Jacobson S, Zibinsky M, Robles O et al. Tumors establish resistance to immunotherapy by regulating Treg recruitment via CCR4. J Immunother Cancer 2020; 8: 764.

52 Newman AM, Steen CB, Liu CL, Gentles AJ, Chaudhuri AA, Scherer F et al. Determining cell type abundance and expression from bulk tissues with digital cytometry. Nat Biotechnol 2019; 37: 773–782.

53 Chen DS, Mellman I. Elements of cancer immunity and the cancer-immune set point. Nature 2017; 541: 321–330.

54 Nicolas-Boluda A, Vaquero J, Vimeux L, Guilbert T, Barrin S, Kantari-Mimoun C et al. Tumor stiffening reversion through collagen crosslinking inhibition improves T cell migration and anti-PD-1 treatment. Elife 2021; 10. doi:10.7554/ELIFE.58688.

55 Sadtler K, Estrellas K, Allen BW, Wolf MT, Fan H, Tam AJ et al. Developing a pro-regenerative biomaterial scaffold microenvironment requires T helper 2 cells. Science (1979) 2016; 352: 366–370.

56 Perez-Penco M, Weis-Banke SE, Schina A, Siersbæk M, Hübbe ML, Jørgensen MA et al. TGFβ-derived immune modulatory vaccine: targeting the immunosuppressive and fibrotic tumor microenvironment in a murine model of pancreatic cancer. J Immunother Cancer 2022; 10. doi:10.1136/JITC-2022-005491.

57 Tian C, Huang Y, Clauser KR, Rickelt S, Lau AN, Carr SA et al. Suppression of pancreatic ductal adenocarcinoma growth and metastasis by fibrillar collagens produced selectively by tumor cells. Nat Commun 2021; 12. doi:10.1038/S41467-021-22490-9.

58 Tian C, Clauser KR, Öhlund D, Rickelt S, Huang Y, Gupta M et al. Proteomic analyses of ECM during pancreatic ductal adenocarcinoma progression reveal different contributions by tumor and stromal cells. Proc Natl Acad Sci U S A 2019; 116: 19609–19618.

59 Bankhead P, Loughrey MB, Fernández JA, Dombrowski Y, McArt DG, Dunne PD et al. QuPath: Open source software for digital pathology image analysis. Sci Rep 2017; 7. doi:10.1038/S41598-017-17204-5.

60 Jensen C, Madsen DH, Hansen M, Schmidt H, Svane IM, Karsdal MA et al. Non-invasive biomarkers derived from the extracellular matrix associate with response to immune checkpoint blockade (anti-CTLA-4) in metastatic melanoma patients. J Immunother Cancer 2018; 6. doi:10.1186/S40425-018-0474-Z.

61 Leeming DJ, He Y, Veidal SS, Nguyen QHT, Larsen D V., Koizumi M et al. A novel marker for assessment of liver matrix remodeling: an enzyme-linked immunosorbent assay (ELISA) detecting a MMP generated type I collagen neo-epitope (C1M). Biomarkers 2011; 16: 616–628.

