## Supplementary data for "Collagen type I promotes pancreatic tumor growth and limits immune cell infiltration"

**Table S1.** List of antibodies

| | Antibody | Fluorochrome | Company | Catalog number |
|---|---|---|---|---|
| Panel 1 | ZombieAqua | | Biolegend | 423101 |
| | Anti-CD45 | PE-Cy7 | Biolegend | 103114 |
| | Anti-CD31 | FITC | Biolegend | 102405 |
| | Biotinylated anti-FAP | | R&D systems | BAF3715 |
| | Streptavidin | APC | Biolegend | 405207 |
| | Anti-PDGFRα | PE | Biolegend | 135905 |
| Panel 2 (myeloid cells) | ZombieAqua | | Biolegend | 423101 |
| | Anti-CD11b | PE-Cy7 | Biolegend | 101215 |
| | Anti-F4/80 | APC | Biolegend | 123116 |
| | Anti-MR | PE | Biolegend | 141705 |
| | Anti-CD11c | FITC | Biolegend | 117306 |
| | Anti-CD103 | BV421 | Biolegend | 121422 |
| | Anti-Ly-6G | APC-Cy7 | Biolegend | 127623 |
| | Anti-Ly-6C | PerCP-Cy5.5 | Biolegend | 128011 |
| Panel 3 (lymphoid cells) | ZombieAqua | | Biolegend | 423101 |
| | Anti-CD3 | FITC | Biolegend | 100204 |
| | Anti-CD4 | BV421 | Biolegend | 100438 |
| | Anti-CD8 | APC | Biolegend | 100712 |
| | Anti-PD1 | PE | Biolegend | 109103 |
| | Anti-CD19 | APC-Cy7 | Biolegend | 108724 |
| | Anti-NK1.1 | PE-Cy7 | Biolegend | 115520 |

**Table S2.** Primer sequences for qRT-PCR

| | Forward | Reverse |
|---|---|---|
| *Actb* | ACTGTCGAGTCGCGTCCA | ATCCATGGCGAACTGGTGG |
| *Col1a1* | TGGTACAGCACTTTACAGCGCACA | TTACTCGGCCTGGGTCACTTCTTT |
| *Cd4* | TTTGCAGAGGAAAACGGGTG | AGAGTCAGAGTCAGGTTGCC |
| *Cd8* | GGATTGGACTTCGCCTGTGA | CTTTCGGCTCCTGTGGTAGC |
| *Nk1.1* | GCTGTGCTGGGCTCATCCT | TTGATGGTTTTTGTACTAAGACTCGCA |
| *Il10* | GCATGGCCCAGAAATCAAGG | GAGAAATCGATGACAGCGCC |
| *Tgfb1* | CTGCTGACCCCCACTGATAC | AGCCCTGTATTCCGTCTCCT |
| *Foxp3* | ATCCATGTGCAAGGAGAGCAG | TTGCCGGGAGCATATACCAGG |
| *Tnfrsf9* | GTCTGTGCTTAAGACCGGGA | GTCTTCTTAAATGCTGGTCCTCC |
| *Pd1* | AACCAGAAGGCCGGTTTCAA | TCCTCTGGCCTCTGACATACT |
| *Tnfa* | CCCACGTCGTAGCAAACCA | CTTTGAGATCCATGCCGTTGG |
| *Ifng* | CGGCACAGTCATTGAAAGCC | TGTCACCATCCTTTTGCCAGT |
| *Gzmb* | AAAGGCAGGGGAGATCATCG | AGGCTGCTGATCCTTGATCG |
| *Cd274* | TCACTTGCTACGGGCGTTT | CCCAGTACACCACTAACGCA |

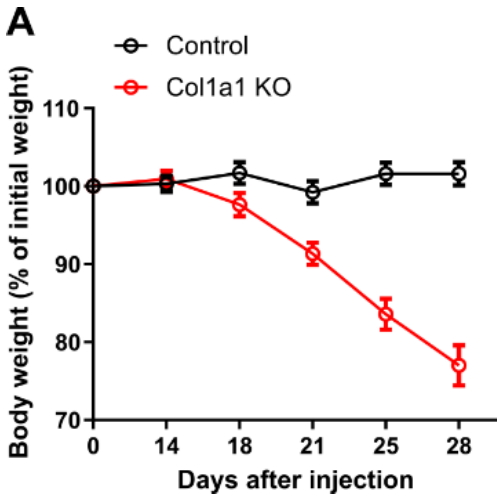


**Figure S1. Weight loss.** Pan02 cells subcutaneously injected in the flank of Col1a1 KO mice or wildtype littermates. Tamoxifen was administered starting when tumors became palpable. The body weight of the mice was measured three times per week until the end of the experiment.

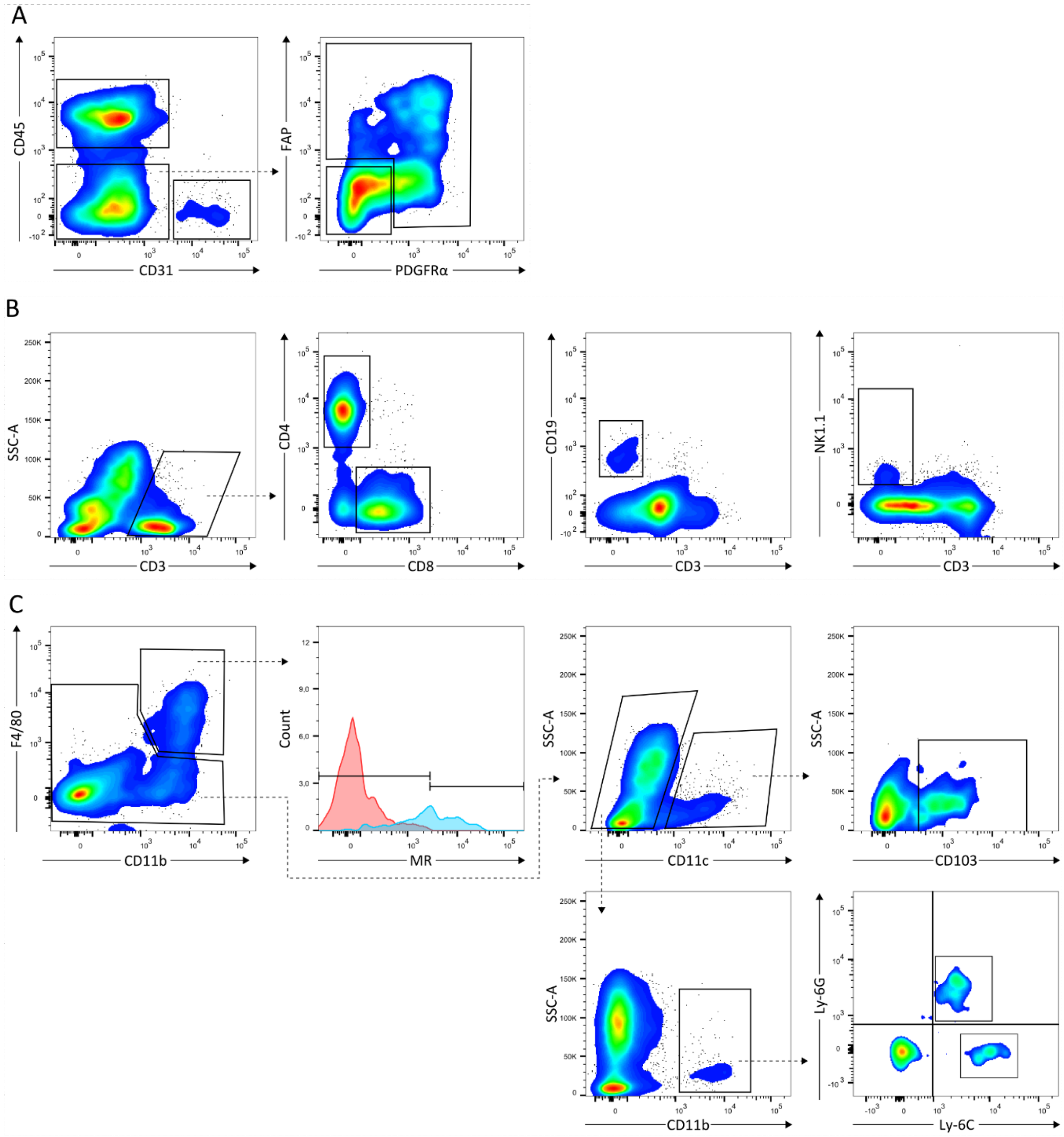


**Figure S2. Gating strategies.** Representative density plots or histograms showing gating strategies from flow cytometry analysis. **(A)** After exclusion of debris, doublets, and dead cells (Zombie aqua-negative), $CD45^{+}$ leukocytes and $CD31^{+}$ endothelial cells were identified. Within the negative population, CAFs were identified based on their expression of FAP and/or PDGFRα. Cancer cells were defined as being negative for all markers (CD45, CD31, FAP, and PDGFRα). **(B)** After exclusion of debris, doublets, and dead cells (Zombie aqua-negative), $CD3^{+}$ T cells were identified. Within the $CD3^{+}$ population, $CD4^{+}$ and $CD8^{+}$ T cells were identified. B cells were identified based on $CD19^{+}$ expression and $CD3^{-}$ expression. NK cells were identified based on their $NK1.1^{+}$ expression and $CD3^{-}$ expression. **(C)** After exclusion of debris, doublets, and dead cells (Zombie aqua-negative), macrophages were identified based on $CD11b^{+}$ and $F4/80^{+}$ expression. Expression of MR on macrophages was measured (representative sample in blue, isotype control in red). Within the $F4/80^{-}$ population, DCs were identified based on $CD11c^{+}$ expression. Within the DC population, $CD103^{+}$ could be identified. In the $CD11b^{+}CD11c^{-}$ population, mMDSCs were defined as $Ly\text{-}6C^{+}Ly\text{-}6G^{-}$ and gMDSCs defined as being $Ly\text{-}6C^{+}Ly\text{-}6G^{+}$.

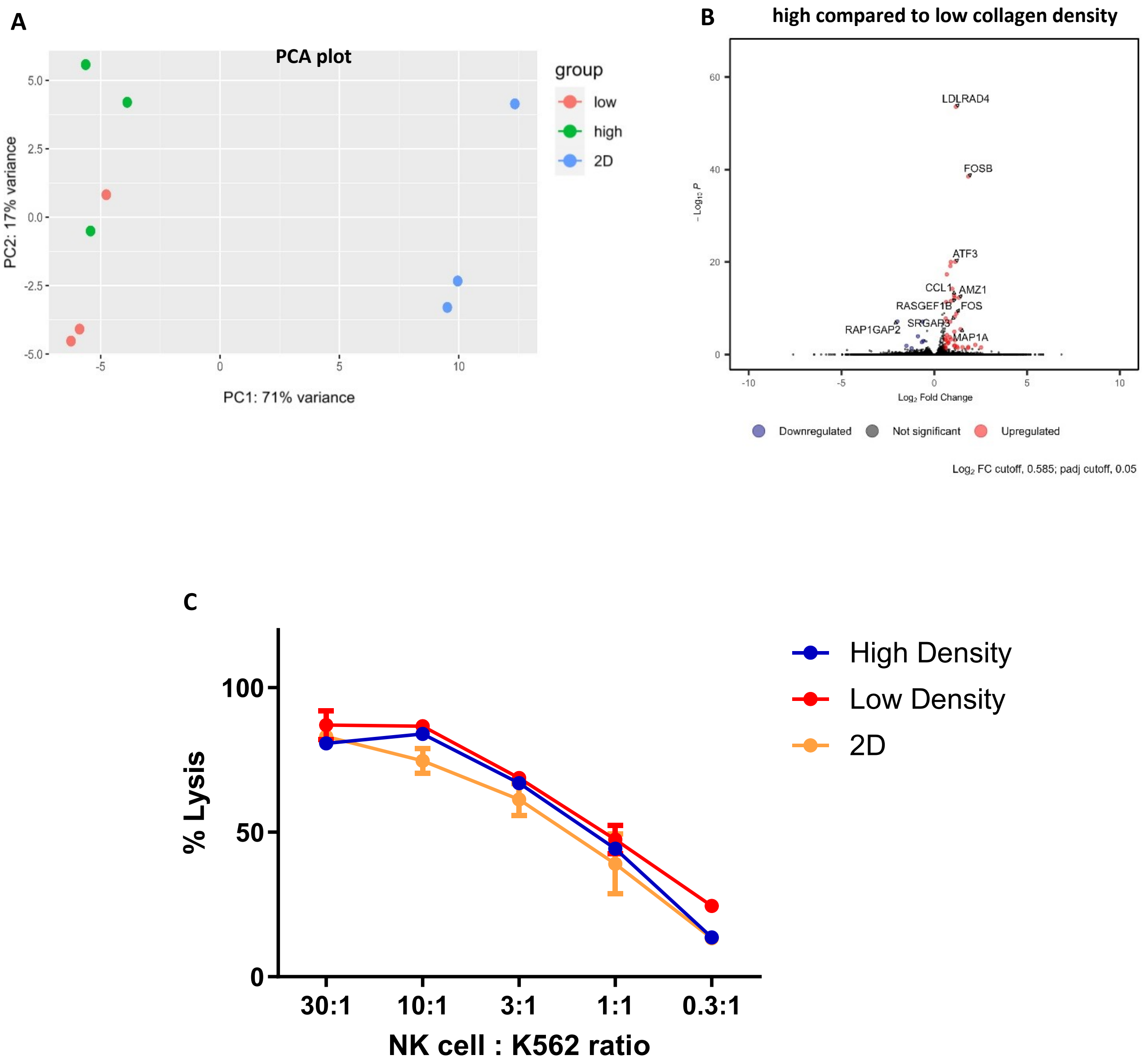


**Figure S3. NK cell culture in different collagen densities. (A-B)** NK cells were analyzed by RNAseq after culture in collagen type I gels of high or low density or on regular tissue culture plastic. (A) PCA plot showing the separation of all samples. **(B)** Volcano plot comparing the transcriptome of NK cells cultured in high collagen density compared to low collagen density. **(C)** After culture of NK cells under different conditions, cells were extracted and their cytotoxic activity was analyzed by co-culture with Chromium-51 pulsed K562 target cells and measuring chromium release.
